# Constructing neural networks with pre-specified dynamics

**DOI:** 10.1101/2023.06.19.545607

**Authors:** Camilo J. Mininni, B. Silvano Zanutto

**Affiliations:** Instituto de Biología y Medicina Experimental, Consejo Nacional de Investigaciones Científicas y Técnicas, Argentina; Instituto de Ingeniería Biomédica, Universidad de Buenos Aires, Argentina

**Keywords:** neural networks, brain dynamics, model fitting

## Abstract

A main goal in neuroscience is to understand the computations carried out by neural populations that give animals their cognitive skills. Neural network models allow to formulate explicit hypothesis regarding the algorithms instantiated in the dynamic of a neural population, its firing statistics, and the underlying connectivity. Neural networks can be defined by a small set of parameters, carefully chosen to procure specific capabilities, or by a large set of free parameters, fitted with optimization algorithms that minimize a given loss function. In this work we alternatively propose a method to make a detailed adjustment of the network dynamic and firing statistic to better answer questions that link dynamic, structure and function. Our algorithm – termed generalized Firing-to-Parameter (gFTP) – provides a way to construct binary recurrent neural networks whose dynamic strictly follows a user pre-specified transition graph that details the transitions between population firing states triggered by stimulus presentations. Our main contribution is a procedure that detects when a transition graph is not realizable in terms of a neural network, and makes the necessary modifications in order to obtain a new transition graph that is realizable and preserves all the information encoded in the transitions of the original graph. With a realizable transition graph, gFTP assigns values to the network firing states associated with each node in the graph, and finds the synaptic weight matrices by solving a set of linear separation problems. We test gFTP performance by constructing networks with random dynamics, continuous attractor-like dynamics that encode position in 2-dimensional space, and discrete attractor dynamics. We then show how gFTP can be employed as a tool to explore the broad dependencies between structure and function, and the specific dependencies subserving the algorithms instantiated in the network activity.

## 1 Introduction

Neural network models play a crucial role in neuroscience, as they enable the formulation of explicit hypotheses that establish connections between behaviuor and neurophysiology. Networks can be constructed from the bottom up, taking experimental evidence to define the neuron input-output mapping and their connectivity. Then, the emergent properties of the system can be studied, expecting that they recapitulate experimental observations not used as model hypothesis. Conversely, a normative approach is also possible, in which we start from the function the network is proposed to have, together with a few assumptions about network connectivity and neuron activation dynamics, and then we fit the free parameters of the model to obtain the best task performance, as measured by a loss function. This approach has increasingly gained momentum due to advances in deep learning that have provided researchers with new tools and hardware to fit increasingly complex models to increasingly complex tasks [1]. The models obtained have reproduced experimental observations regarding the nature of neural coding, distributed representation and population dynamics [2, 3, 4]. Modelling approaches of this kind stem from a paradigm that proposes to understand neural computation as algorithms instantiated in the low dimensional dynamic of large neural populations [5, 6]. With neural recording technology allowing more simultaneous measurements [7] fitting neural networks with many neurons, capable of complex dynamics is going to be more and more necessary. Nonetheless, network optimization is a complex endeavour. The way fitted networks solve the task, and how they represent information depend on the choosing of all hyperparameters, making it difficult to draw conclusions about how each hypothesis impacts on network behaviour. Moreover, a fitting algorithm may converge preferentially to a specific family of solutions, giving no information about how many other networks there are that can solve the task in qualitatively different ways. Neither the inability to achieve low error in a given task rules out the proposed hypotheses as candidates to explain a phenomenon. Furthermore, there are many synaptic weight configurations that lead to the same network output, making it difficult to relate network function with network connectivity. Training is usually easier if networks have many neurons (with many parameters), hence there is a bias towards choosing networks with more neurons than the minimum required, making the number of weight configurations even larger. However, some of these shortcomings can be handle by leveraging the particular goals of modelling in neuroscience. First, we expect our network to solve tasks alike the ones solved by animals in behavioural experiments. These tasks are usually easy to solve, in the sense that we already know how to solve them. We can even enumerate more than one algorithm that solves the same task. Second, we usually have some hypotheses about how the modelled system represents and processes information. For instance, we may have evidence that neural populations in the prefrontal cortex handle working memory demands by instantiating discrete attractors that encode relevant stimuli[8, 9], or that networks in the entorhinal cortex encode position in a 2D continuous attractor[10, 11, 12]. Therefore, it would be of interest to have a network that solves a task through a given *a priori* dynamic, and then study how the underlying connectivity relates to function. However, specifying the dynamic through minimization of a loss function by means of an optimization algorithm could be very hard to accomplish. In a previous work we proposed an algorithm that constructed networks by solving a system of linear equations [13]. In this algorithm there was one equation for each transition in the transition graph, the coefficients were the firings of each neuron in the source state, and the variables were the membrane potentials in the target state. The critical aspect of the algorithm was a way to define the membrane potentials such that the system of equations had an exact solution. We successfully employed the algorithm in a particular task -a sequence memory task-, but how to generalize the algorithm to arbitrary transition graphs with any number of stimuli remained an open question. In the present work we described a much more general algorithm: the generalized Firing-to-Parameter (gFTP) algorithm. This new version takes a transition graph as input, decides if there is a neural network capable of following the transition graph exactly, and if not, expands the original graph to construct a new graph that fulfil two conditions: it is realizable in terms of a neural network, and it retains all of the information present in the original graph. Then, we define the network firing states associated with each node in the graph, in such a way that linear combinations induced by stimuli are satisfied. Finally, weight matrices are found with the accelerated perceptron algorithm [14].

In the following, we start by giving a detailed description of the rationale behind the algorithm, and evaluate its performance on: random transition graphs, graphs instantiating a discretized version of a 2-dimensional continuous attractor that encodes position, and graphs instantiating a discrete attractors dynamic. Then, we show how gFTP can be employed in combination with an optimization algorithm as an exploratory tool to disentangle the multiple dependencies between structure and function. Finally, we analyse the underlying differences in dynamic and connectivity subserving two different algorithms that solve the same behavioural task.

## 2 The Generalized Firing to Parameter Algorithm

### Network model and consistency conditions

We will consider recurrent neural networks composed of binary neurons (Fig. 1a). Neurons receive inputs from sensory neurons and from all other neurons in the network. Sensory neurons encode stimuli in a one-hot fashion. The network activity at iteration *k* is dictated by:

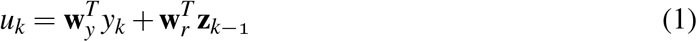

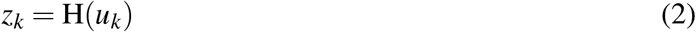

where *u* is the preactivation of the neuron, **y** is the vector that collects the activations of sensory neurons (one neuron per stimulus), **z** is the vector that collects the activations of neurons in the recurrent network and H the Heaviside function. Vectors **w**_*y*_ and **w**_*r*_ are the synaptic weights of the inputs coming from the sensory neurons and neuron in the recurrent network respectively, for a given neuron in the network. Conversely, matrices **W**_*y*_ and **W**_*r*_ collect in their rows the synaptic weight vectors 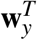 and 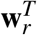 of all neurons in the recurrent network (Fig. 1f).

**Figure 1.**
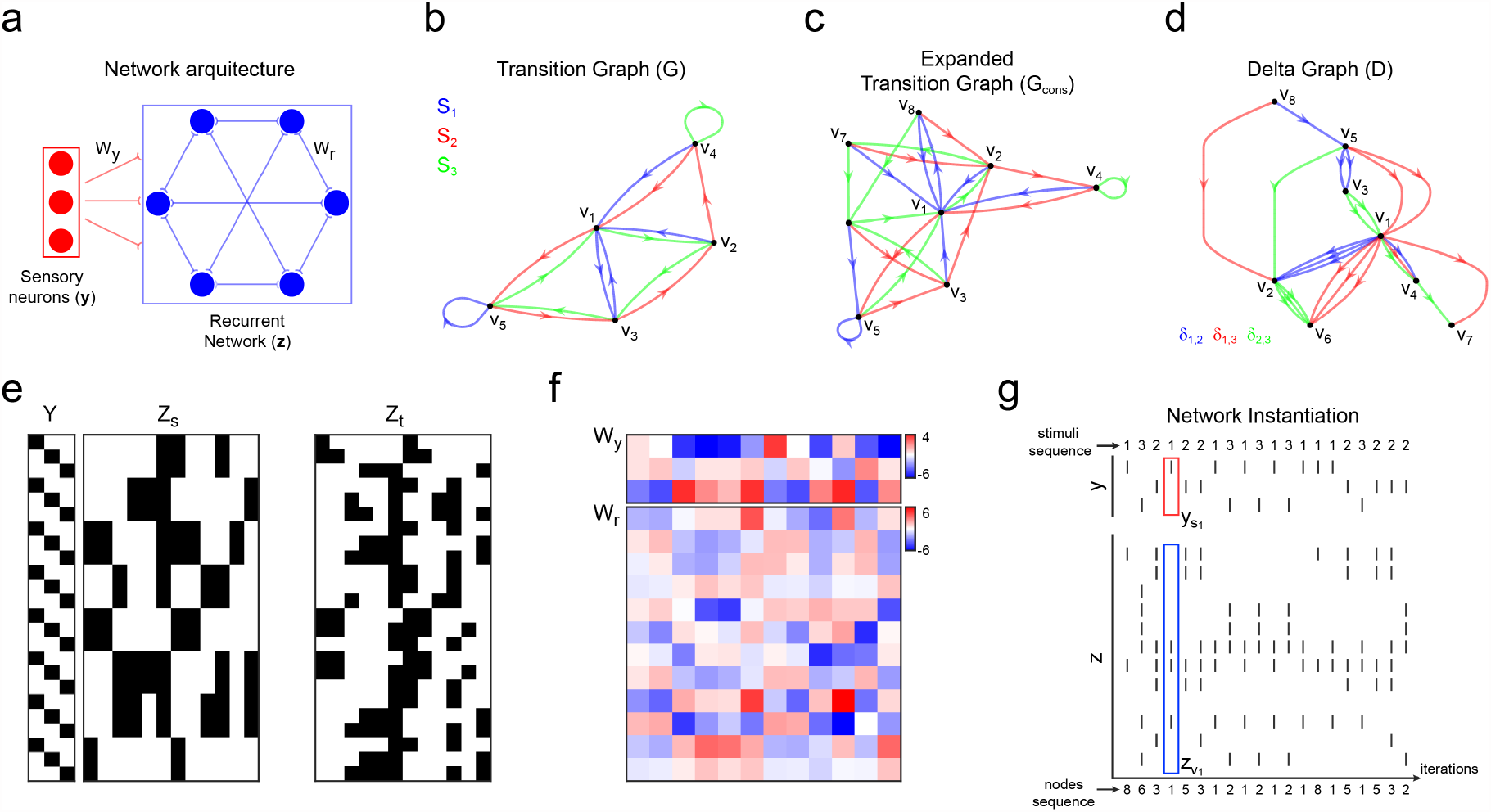
Recurrent neural networks that follow a pre-specified transition graph. **(a)** Recurrent networks of binary neurons receive connections from neurons that encode stimuli in a one-hot fashion. **(b)** An example of transition graph that defines the desired network dynamic. Each node represents a different neural population state in the recurrent network. Directed arcs depict transitions from source to target states that are triggered by different stimuli, encoded by the sensory neurons and shown with arrows of different colors. **(c)** Graph en (b) was expanded until no inconsistencies remained. **(d)** Graph *D* computed from the expanded graph in (c). Arc color represents the arc label (delta value, one for each stimuli pair). The graph shows no cycles, confirming that the expanded graph is now realizable in terms of a neural network. **(e)** Matrices **Y, Z**_*s*_, and **Z**_*t*_, constructed by backtracking through the neurons activations that differentiate all nodes and comply with delta constraints. **(f)** Synaptic weight matrix found through the accelerated perceptron algorithm, with matrix [**Y, Z**_*s*_] collecting samples to classify, and matrix **Z**_*t*_ collecting the binary classes. **(g)** Raster plot showing the activity of a neural network constructed to follow expanded graph in (c) during presentation of a sequence of random stimuli.

We want to find the synaptic weight matrices **W**_*y*_, **W**_*r*_ that instantiates a network that unfolds some target dynamic. Network dynamics can be specified in the form of a labeled multi digraph *G* = (*V, K, S, f*_*s*_, *f*_*t*_, *l*_*G*_), where *V* is the set of nodes 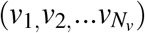, *K* ⊂ ℕ is the set of transitions specified (the arcs in the graph), *S* ⊂ ℕ the set of stimuli (which act as labels) the network can have as inputs, *f*_*s*_ the function that takes a transition and gives the source node, *f*_*t*_ the function that takes a transition and gives a target node, and *l*_*G*_, the function that takes a transition and gives the label (the stimulus that triggers that transition) (Fig. 1b). Conversely, a matrix **G** can be constructed, such that row *k* is (*l*_*G*_(*k*), *f*_*s*_(*k*), *f*_*t*_(*k*)). When unambiguous, we will use subindexes to indicate that a given state vector is associated with another vector in a certain way (for example, that **z**_*v*_ is the **z** that encodes node *v*).

We define matrices **Y, Z**_*s*_ and **Z**_*t*_ such that each column collects the activations of the sensory neurons (**Y**), and the activation of the recurrent network in a source state (**Z**_*s*_) and in a target state (**Z**_*t*_), for each transition/column *k* (i.e. **Y**(:, *k*) = **y**_**G**(*k*,1)_, **Z**_*s*_(:, *k*) = **z**_**G**(*k*,2)_) and **Z**_*t*_(:, *k*) = **z**_**G**(*k*,3)_,(Fig. 1e)). A network that instantiates graph *G* must satisfy:

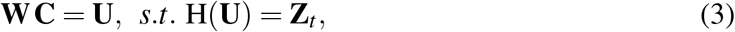

with

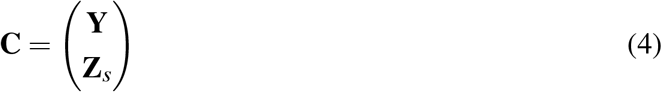

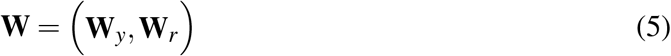

This is a problem of linear separability. Matrix **W** exists iff. *rank*(**C**) = *rank*((**C**^*T*^, **U**^*T*^)), (i.e. the augmented matrix (**C**^*T*^, **U**^*T*^) and matrix **C** present the same linear combinations), and the signs of values in **U** are consistent with **Z**_*t*_. If **Y, Z**_*s*_ and **Z**_*t*_ are chosen properly, then a solution for equation 3 exists and can be found with recently proposed accelerated versions of the perceptron algorithm. Finding suitable matrices is a hard combinatorial problem. However, we can leverage some regularities to reduce its complexity. For a start, we must first find if such **Z**_*s*_, **Z**_*t*_ exist in the first place, since it could be the case that, for some transition graphs, the interdependencies between source and target nodes preclude its instantiation in the shape of any neural network. In this sense, we must find out if the graph is realizable, and if not, we must find a way to turn the graph into a realizable graph, without loosing the critical aspects of the target dynamic.

There is a set of linear combinations that directly stems from the transition graph and can be detected and corrected if needed. Let’s consider two transitions that start from the same source node *v*, but are triggered by different stimuli:

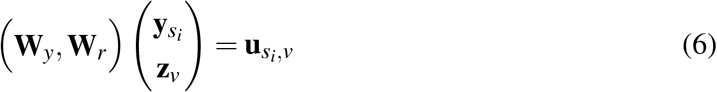

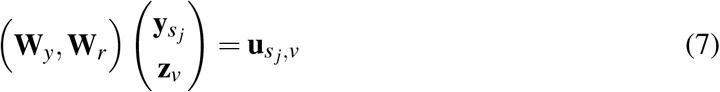

where 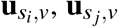 are the preactivations after the network receives stimulus *s*_*i*_ or *s* _*j*_, starting from the same node *v*. Subtracting both sides of the equations we get:

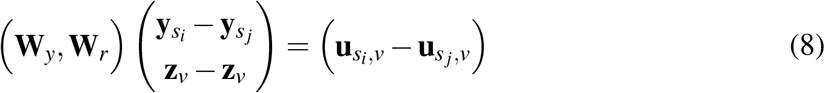

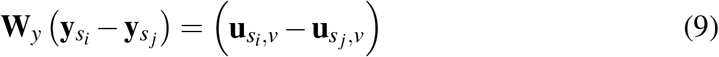

Since both transitions start from the same node, the inputs that a neuron receives that come from the other neurons in the recurrent network cancel, and the difference in **u** only depends on the input from the stimuli. Vector 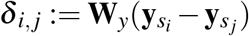 is a constant that does not depend on the source node *v*. It follows that vector 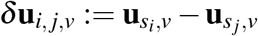 does not depend on *v* either, since it must be equal to *δ* _*i, j*_ for all *v*. For a given neuron *n*:

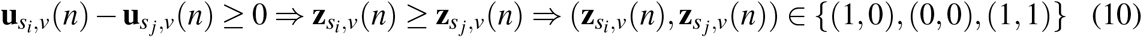

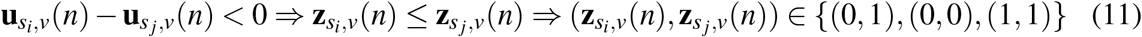

Therefore, all pairs of **z** that are reached from a given pair of stimuli from the same source are restricted on the values that can jointly adopt. Not taking this fact into consideration will lead to **Z**_*s*_ and **Z**_*t*_ that are not realizable in any neural network. Moreover, it could be the case that the graph itself does not allow *any* **Z**_*s*_, **Z**_*t*_ to be realizable. Consider for example the graph in Fig. 2a:

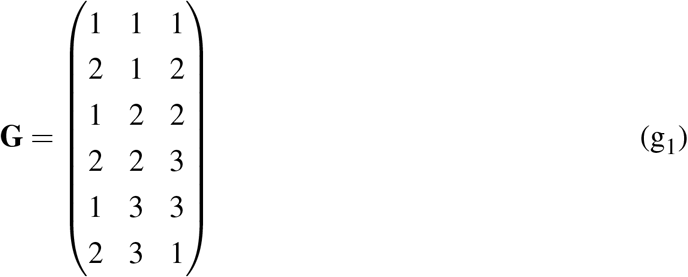

If a neuron differentiates *v*_1_ from *v*_2_, by firing when the network is in state 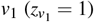 and not firing in state 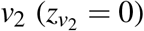, then *δ*_1,2_ *>* 0. This in turn means that the neuron must not fire in state 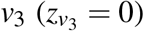, because if it fired then *δ*_*i, j*_ *<* 0, leading to a contradiction to the value of delta defined by transitions 1 and 2. We say that the value of zero assigned to *v*_2_ “propagates” to *v*_3_. This value propagates from *v*_3_ to *v*_1_ too 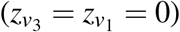, where we started. But this is in contradiction with the original value for 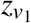. The inconsistency is circumvented only when *v*_1_ and *v*_2_ are not differentiated by any neuron, but in this case there would be only two network states instead of three. Therefore, the graph is unrealizable as it is. However, the graph can be modified to make it realizable. Let’s consider the modified graph in Fig. 2b:

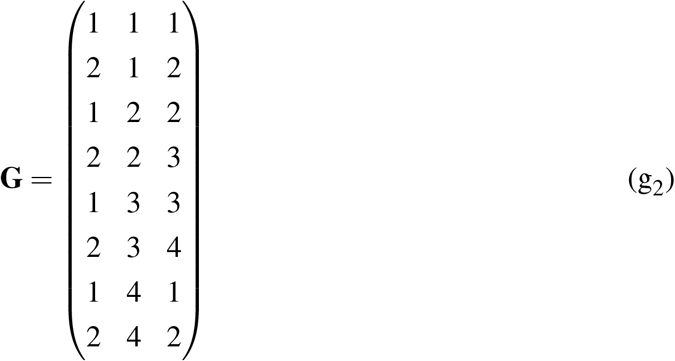

We have replaced *v*_1_ in transition 6 by a new node (*v*_4_), and added transitions for this new node as source, equal to the transitions of the node it replaced (transitions with *v*_1_ as a source node). Since the new node does not appear anywhere else, a vector **z** can be assign according to any *δ* _*i, j*_, without possibility that it contradicts a previous assignment, as it occurred in graph (g_2_). On the other hand, this new node encodes the same information than the replaced node (i.e. the occurrence of stimulus *s*_2_ starting from *v*_3_), and leads to the same transitions too (going to *v*_1_ through *s*_1_ and *v*_2_ through *s*_2_). Therefore, the graph has been altered and it is now realizable, and all the information encoded in the network states is preserved. The new node is a twin of the node it replaces, in the sense that it leads to the same targets through the same stimuli. Note that, if we had a graph of the type:

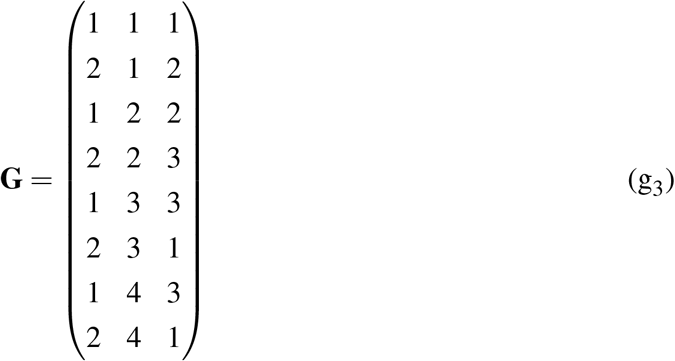

we would need to replace node *v*_1_ in two transitions (transition 6 and 8) in order to attain consistency. We will say that a transition graph is *delta consistent* if there are **Z**_*s*_, **Z**_*t*_ and **U** such that relations (10-11) are satisfied. If this is not the case (because differentiating some nodes leads to a conflict with relations (10-11)) then the graph is termed *delta inconsistent*. We will name “expansion” to the action of replacing a target node by a new node at a set of conflicting transitions, as we did with graph (g_1_). Each expansion removes one inconsistency.

**Figure 2.**
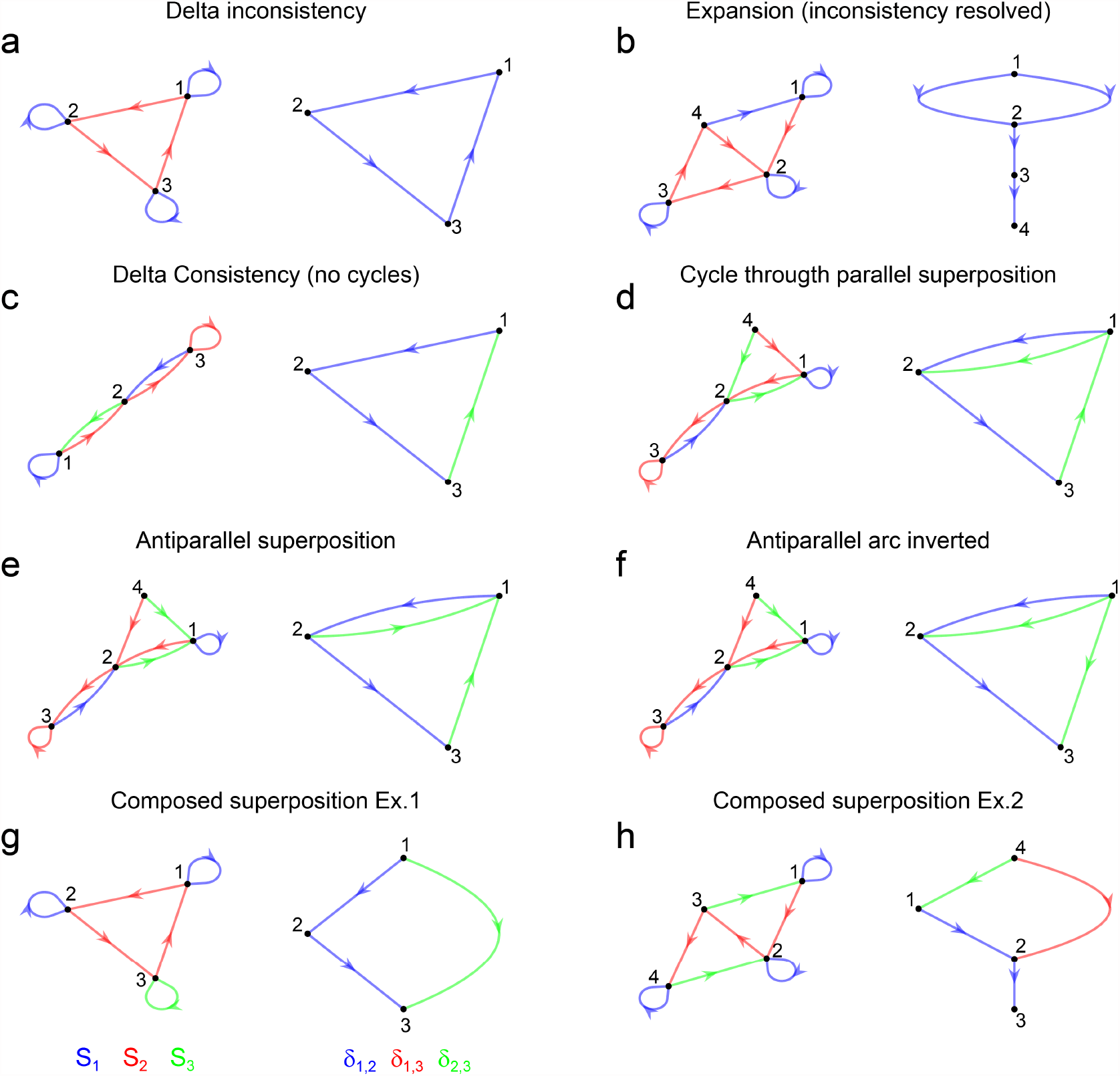
Example graphs *G* (left of each panel) and their associated graph *D* (right of each panel). **(a)** A cycle of 3 nodes. If a neuron is active in one node and not active in the other node, we arrive at a contradiction. This is evidenced by the presence of a cycle in graph *D*. (b) Inconsistency in (a) is resolved by adding node *v*_4_ with same targets than *v*_1_ (node *v*_1_ is expanded). The associated graph *D* has no cycles. **(c)** A graph with 3 nodes and 3 stimuli. Graph *D* has no cycles because it is not possible to flow from a blue arc to a green arc in graph *D* (there is no superposition of deltas). **(d)** Blue and green arcs in graph *D* are superimposed, putting them in the same traversable set. Hence, there is a cycle. **(e)** Blue and green arcs in graph *D* are superimposed but they have opposite directions. These deltas are linked, and arcs labeled with one of these deltas must be inverted. **(f)** Same graph *D* as in (e) but green arcs have been inverted. All arcs belong to the same traversable set, but no cycles are present. Therefore, graph *G* is left unchanged. **(g)** Arcs blue and green are not directly superimposed but they are linked: defining *δ*_1,2_ or *δ*_2,3_ also defines *δ*_1,3_. **(h)** Composed superposition appears any time two nodes are reachable by more than one path (regardless of arcs direction).

In graph (g_1_) we see that the inconsistency appears because we differentiate *v*_1_ from *v*_2_, defining *δ*_*i, j*_ in the process. Assigned values “propagate” to other nodes, reaching back to one of the nodes we already defined. Therefore, an inconsistency appears because we tried to differentiate two nodes that are part of a certain kind of *cycle*. To make this intuition more precise we define a directed multi digraph *D* = (*V, A,C, g*_*s*_, *g*_*t*_, *l*_*D*_),where *V* is the set of nodes,

*A* ⊂ ℕ is the set of arcs, *C* = {*δ*_*i, j*_ : (*i, j*) *∈ S×S*},the set of labels for the arcs, *g*_*s*_ the function that takes an arc and gives its source node, *g*_*t*_ the function that takes an arc and gives its target node, and *l*_*D*_ the function that takes an arc and gives its label (Fig. 1d). An arc *a* from source node *v*_*i*_ to target node *v* _*j*_ is in *A* if: *v*_*i*_ ≠ *v* _*j*_, *l*_*D*_(*a*) = *δ*_*m,n*_ and exist *k*_1_, *k*_2_, *v*_*s*_ such that *f*_*t*_(*k*_1_) = *v*_*i*_, *f*_*t*_(*k*_2_) = *v* _*j*_, *f*_*s*_(*k*_1_) = *f*_*s*_(*k*_2_) = *v*_*s*_ and *l*_*G*_(*k*_1_) = *m, l*_*G*_(*k*_2_) = *n*. In other words, two nodes are connected by an arc in *D* if in graph *G* the nodes are target nodes of the same source node. The label of the arc is *δ*_*m,n*_, the delta associated with the stimuli that lead to each of the connected nodes. The direction of the arc is given by the sign of *δ*_*m,n*_. Self loops (*g*_*s*_(*a*) = *g*_*t*_(*a*)) are not included in *D* because they represent cases in which *δ* = 0, which does not constrain assignments to *z* in any way.

Graph *D* connects nodes if they are linked by deltas, as we have reasoned above. If this graph has a cycle, then there is at least a pair of nodes that cannot be differentiated without getting a delta inconsistency. Therefore, one way to assure delta consistency is to construct graph *D* and search for cycles. If a cycle is found, then we must break it by breaking any of the arcs that composes it. Since an arc exists when two nodes are target of the same source node, the arc can be broken by expanding one of the nodes of the arc, at all transitions in *G* that allow the arc to exist. The expansion breaks the cycle. We have to repeat the process of search and expansion until no cycles are present. The final graph will be delta consistent, and will hold all the information encoded by the original graph (Fig. 1c, d, g).

Navigating the graph *D* is related to differentiating nodes and propagating *z* values. Two nodes can be connected by 2 or more arcs of different label. Consider the graph in Fig. 2c:

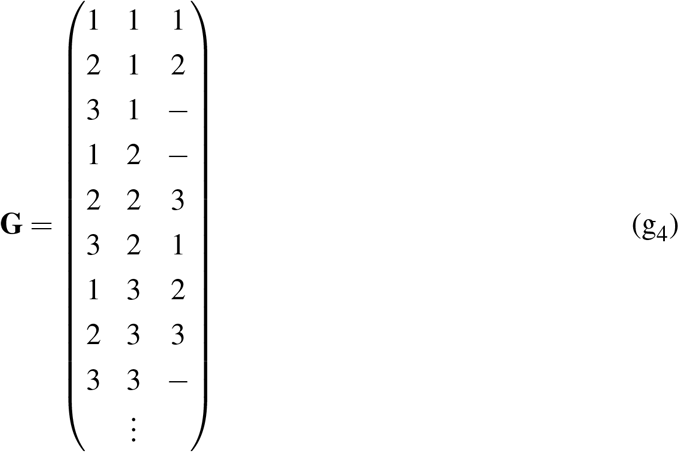

This subset of the graph shows no cycles, and nodes *v*_1_, *v*_2_ and *v*_3_ could be so far differentiated. However in the graph of Fig. 2d:

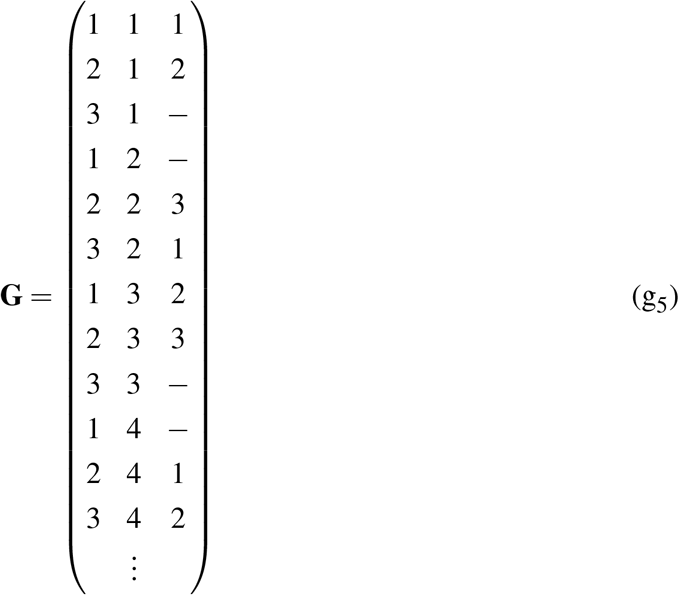

node *v*_1_ and *v*_2_ are connected by two arcs with different labels, meaning that differentiating these two nodes will define two different deltas. Therefore, this graph do present a cycle. *Arc label superposition* links the deltas that label the arcs involved, since defining one delta necessarily defines the other. In turn, arcs with linked deltas can be traversed as if they had the same label. Linked deltas can be represented as a partition *P* of the set of labels *C* (i.e. *P* = {*ℒ*_*i*_}_*i*_, with ∩_*i,j*_(ℒ_*i*_, *ℒ*_*j*_) = Ø, ∪_*i*_ℒ_*i*_=*C*). Labels *δ*_*i, j*_ and *δ*_*m,n*_ are in an element of the partition ℒ_*k*_ if there are arcs *a, b* such that *g*_*s*_(*a*) = *g*_*s*_(*b*) = *v*_*s*_, *g*_*t*_(*a*) = *g*_*t*_(*b*) = *v*_*t*_ and *l*_*D*_(*a*) = *δ*_*i, j*_, *l*_*D*_(*b*) = *δ*_*m,n*_. Then, for each element of the partition, there is one *traversable set* of arcs *V*_*trav*_(ℒ_*i*_) = {(*v*_*s*_, *v*_*t*_) : *l*_*D*_(*a*) *∈ ℒ*_*i*_, *v*_*s*_ = *f*_*s*_(*a*), *v*_*t*_ = *f*_*t*_(*a*), *a* ∈ *A*}. Before detecting any superposition, deltas are only linked with themselves (*a* = *b*), and hence *P* = *C*. As superimposed arcs are detected, deltas are linked by merging the elements of the partitions that contain the linked deltas.

In brief, to find cycles in graph *D* we choose a set ℒ from *P*, and then a source node from *V*_*trav*_(ℒ). Next, we perform depth first search (DFS), traversing the graph through arcs that are members of *V*_*trav*_(ℒ). When a cycle is encountered, a node from the cycle is expanded, graph *G* and *D* are updated and the process is restarted from a new source node in *V*_*trav*_(ℒ). When no cycles are encountered withing one traversable set, a new ℒ is chosen and its associated *V*_*trav*_(ℒ) explored. If all traversable sets were explored and no cycles were found, the process stops and returns the final consistent graph *G*_*cons*_.

Arc superposition can be parallel, like in the example above (superimposed arcs have the same direction), but can also be antiparallel (arcs in opposite direction). In this case, differentiating *v*_1_ from *v*_2_ defines two deltas, but the graph cannot be traversed in the same way as when superposition goes in the same direction. Consider the graph in Fig. 2e:

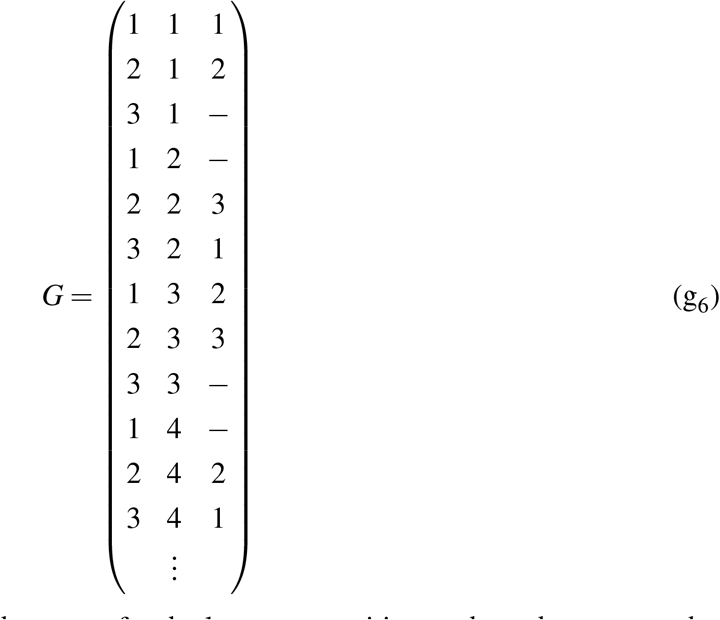

which is equal to the previous graph, except for the last two transitions, where the target nodes where switched. In this case, no inconsistency occurs. Yet, a cycle would be detected if the algorithm so far described is executed. This issue is solved by noting that antiparallel superposition implies that deltas have opposite sign, meaning that the graph should be traversed in opposite direction for arcs of superimposed labels. Hence, for each case of antiparallel superposition we flip the direction of all arcs that share the label with one of the antiparallel superimposed arcs (Fig. 2f), with the precaution of only flipping arcs that weren’t already flipped.

Finally, we can see that the parallel and antiparallel superpositions shown so far are particular examples of a broader definition of superposition. Consider the graph in Fig. 2g:

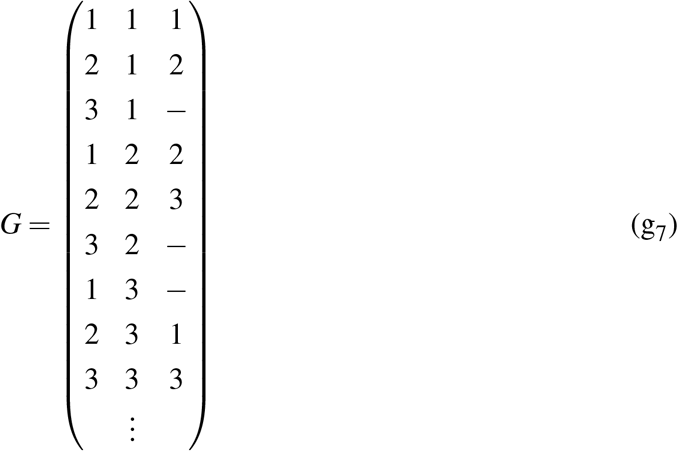

In this case, differentiating *v*_1_ from *v*_2_ defines *δ*_1,2_ but also *δ*_1,3_. In the graph of Fig. 2h:

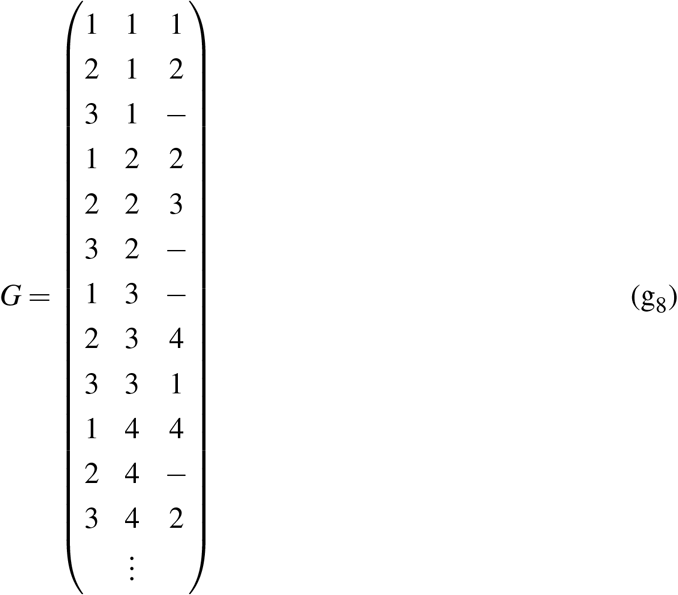

differentiating *v*_1_ from *v*_2_ defines *δ*_1,2_ but also one of the other two deltas: *δ*_4,2_ (parallel to *δ*_1,2_) or *δ*_4,1_ (antiparallel to *δ*_1,2_). It is clear that superposition can occur between two arcs that do not share their source and target nodes. We say it is a *composed superposition*. With this in mind we are able to propose a more general way of detecting arc label superposition. We first define a *complete path* as a path composed of arcs that are within a traversable set, which may or may not be a cycle, but has the property that the first node in the path is never a target node in the traversable set, and the last node is never a source node in the traversable set. Then, if it is possible to go from one node of the complete path to another node of the complete path, traversing through arcs contained in any of the *remaining* traversable sets (without taking their direction into consideration), then one of the labels of the arcs that connect the two nodes is superimposed to the arcs of the complete path. The superposition is parallel, if the superimposed arc was traversed in the direction of the arc, or antiparallel otherwise. Once the superposition is detected, elements of the partition *P* and the associated traversable sets are merged to account for the new linked deltas. Note that this way of detecting composed superposition will also detect simple parallel and antiparallel superpositions as the ones in examples g_5_ and g_6_. In gFTP, we start by finding the partition *P* and traversable sets, as defined above. Then, we find all the complete paths and detect the composed superpositions. Partition *P* and traversable sets are modified according to the presence of parallel and antiparallel superposition. For antiparallel arcs, we invert the direction of all the arcs that share the label with the arcs of the antiparallel pair of arcs. Arcs that were flipped in their direction are not flipped again. Finally, we merge elements of *P* and traversable sets according to the superpositions encountered (see the Appendix for pseudocode of the gFTP algorithm and its subfunctions).

## Construction of Z_*s*_ and Z_*t*_

Matrices **Z**_*s*_ and **Z**_*t*_ are generated in a neuron-by-neuron basis, i.e. the neuron activation states

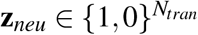 are defined by defining its output after each transition (**z**_*neu*_ is a row in matrix **Z**_*t*_, of number of elements equal to the number of transitions *N*_*tran*_). Finding **z**_*neu*_ is a constrained satisfaction problem, which we solve with a backtracking algorithm, where there is one binary decision for each node, and the constrain to satisfy is delta consistency. Values 1 or 0 are assigned to nodes, and if delta consistency is sustained, we search for nodes that are obliged to have a certain value, either 1 or 0, such that delta consistency is not violated, i.e. assigned values are “propagated” to other values, according to deltas defined so far (for example, if *δ*_*i j*_ = *u*_1_ − *u*_2_ *>* 0 and *z*_1_ = 0, then *z*_2_ must be 0 in order to sustain delta consistency). Value propagation using delta consistency helps in reducing the number of combinations to try. To reduce it further, nodes are explored in decreasing order of their in-degree in graph *G*_*cons*_. This follows from the assumption that nodes that appear more times as targets in *G* have more instances to satisfy a given delta. This should make their assignment more difficult and critical to following assignments, and thus they should be assigned first. Details on our backtracking algorithm for constructing **Z**_*s*_ and **Z**_*t*_ can be found in the *Methods* section.

## Overview of the gFTP algorithm

1. Graph *D* is constructed from graph *G*
2. Arc superpositions are detected and traversable sets are updated
3. Graph *D* is traversed through DFS.
4. If a cycle is found, one of its nodes is expanded
5. Steps 1 to 4 are repeated until no cycles are present
6. Matrices **Z**_*s*_ and **Z**_*t*_ are constructed
7. Matrices **W**_*y*_ and **W**_*r*_ are obtained with the accelerated perceptron algorithm
8. If step 7 fails, new matrices **Z**_*s*_ and **Z**_*t*_ are constructed and concatenated to the previous ones
9. Steps 7 and 8 are repeated until step 7 is successful

## 3 Results

### Algorithm performance

We employed gFTP to construct networks from three kinds of transition graphs: random transition graphs (Fig. 3a), transition graphs that encode position in 2-dimensional space (Fig. 3b), and transition graphs that instantiate discrete attractor dynamics (Fig. 3c) (see details in *Methods*).

**Figure 3.**
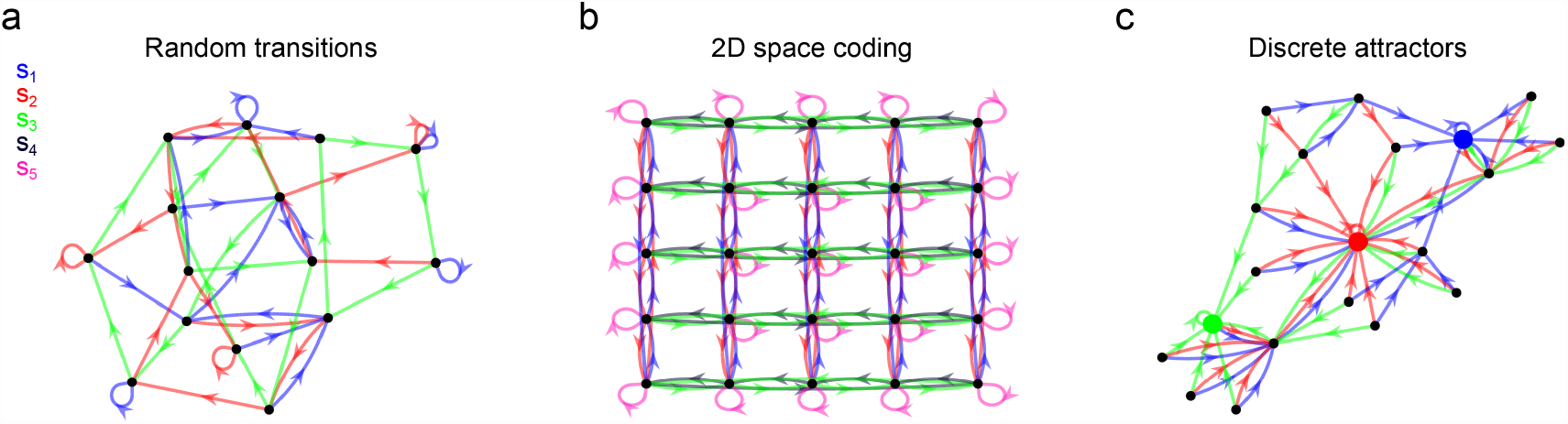
Examples of the transition graphs evaluated (before expansion) **(a)** Random graph with 15 nodes and 3 stimuli. A source node *i* has target nodes taken at random from *i*− 2 to *i* + 2. **(b)** Transition graph of a network that encodes 25 discrete positions in a 2-dimensional square space, of side length = 5. There are five stimuli, associated with displacement to the right (green), left (grey), up (blue), down (red), and a fifth stimulus (pink) associated to stillness. **(c)** Transition graph with 3 nodes acting as attractors (depicted with bigger markers). Each stimulus leads to one specific attractor through a short-distance path. Graph shown is before completing nodes without source.

We measured the wall-clock execution time of the algorithm for each kind of transition graph. The complexity of the graph was adjusted by setting the number of nodes or stimuli in random graphs, the side-length of the arena in 2-dimensional space encoding graphs, or the number of nodes in the case of discrete attractor graphs. We separately measured the time required to obtain a consistent graph (consistency time, Fig. 4) and the time to construct **Z**_*s*_, **Z**_**t**_ and **W** matrices (construction time, Fig. 5). Consistency time was well fitted by a power function and not an exponential (Fig. 6), suggesting polynomial time complexity to obtain a realizable graph from a starting random graph. The number of nodes in the consistent graphs was higher than in the initial graphs (except for the discrete attractors graphs), and the relation between these two numbers was almost linear. Execution times differed considerately between graph types. Lower consistency times were found for discrete attractor graphs, followed by random graphs with 3 stimuli, random graphs with 6 stimuli, and 2-dimensional space coding graphs. Neither consistency nor construction times were fully explained by the number of nodes in the final graphs, suggesting that the shape of the graphs was an important factor in determining the number of steps necessary for making the graph consistent, and for constructing the network. The number of neurons *N*_*neu*_ in the constructed network grew linearly with graph size, and was close to the number of nodes in the final graph. This implies that, in most cases, 1 neuron per node was enough, since gFTP adds neurons to the network in multiples of the number of nodes in the consistent graph.

**Figure 4.**
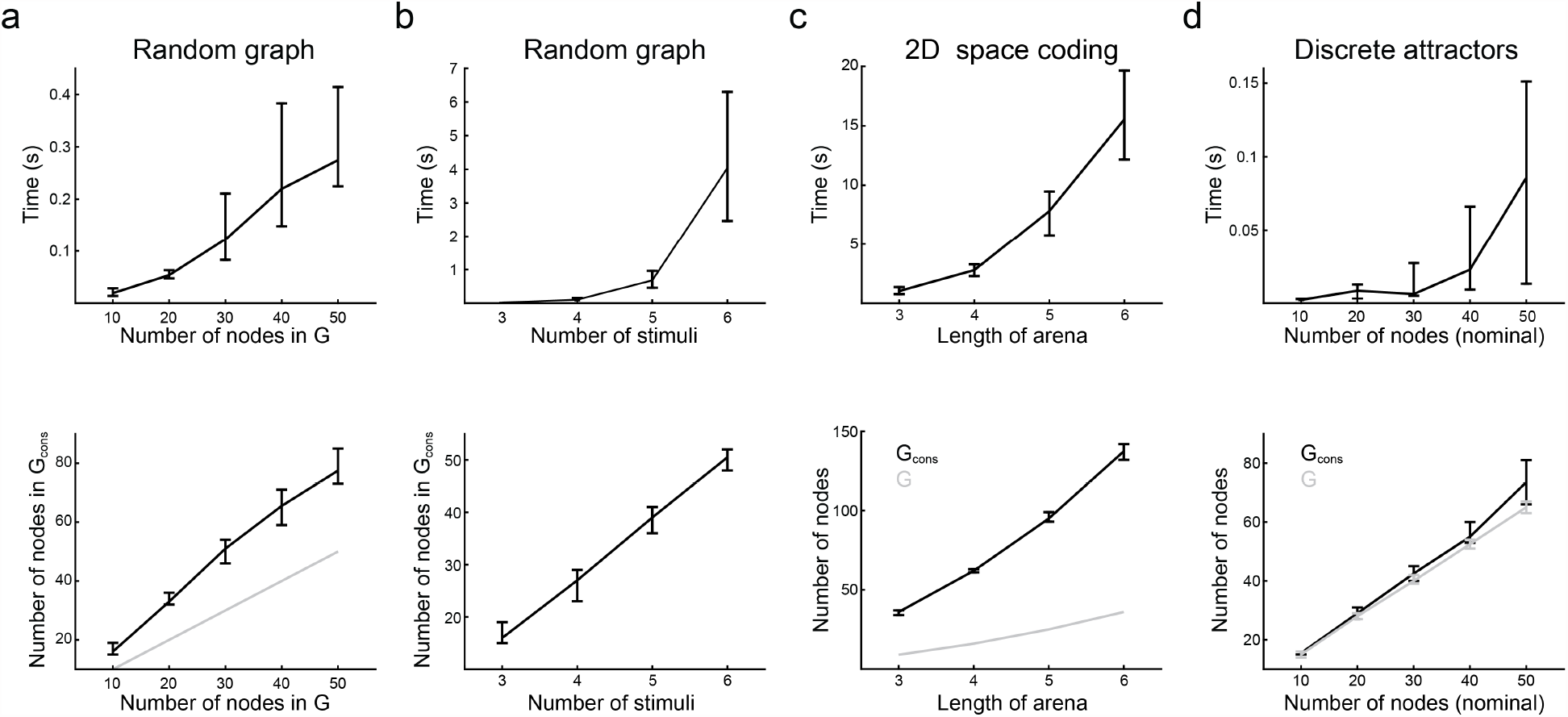
Time performance during consistency assessment. **(a-d)** Execution time until a consistent graph is obtained (upper row), and number of nodes (lower row), for the three types of transition graphs tested. *G*: initial graph, *G*_*cons*_: consistent graph obtained after successive cycle detection and node expansion. **(a)** Random graphs with 3 stimuli and increasing number of nodes. The grey line is the identity function. **(b)** Random graphs with 10 nodes and increasing number of stimuli. **(c)** 2-dimensional space coding graphs, with 5 stimuli and (length of arena)^2^ number of nodes (grey line). **(d)** Discrete attractor graphs, with 3 attractors, one stimulus per attractor that leads to it, and an increasing number of nodes. The *x* axis shows (nominal) number of nodes in the graph (before adding new source nodes to sourceless nodes). The *y* axis of the lower panel shows the final number of nodes in *G* after correcting for sourceless nodes, and the final number of nodes in *G*_*cons*_. Median *±* [25th, 75th] percentiles are shown, computed over 30 networks for each condition.

**Figure 5.**
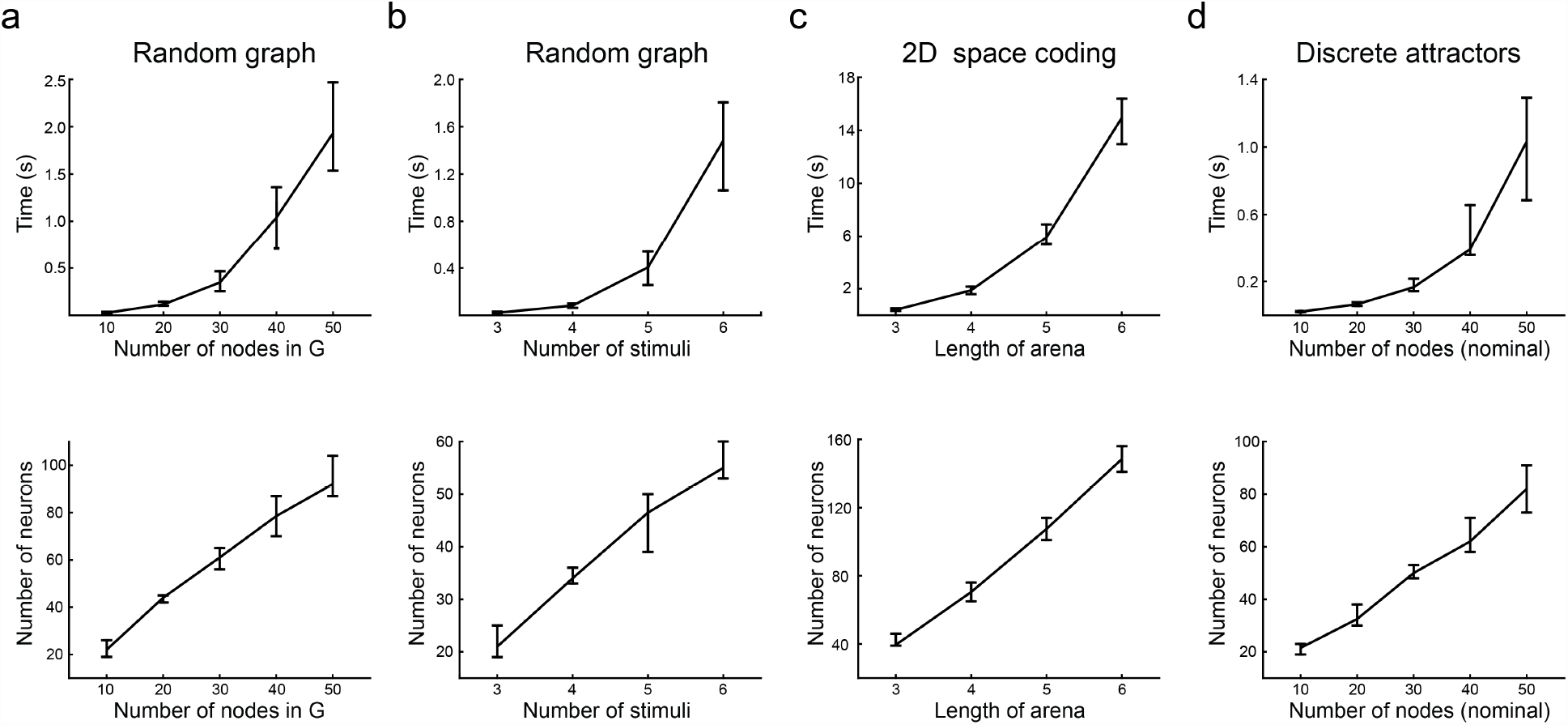
Time performance during network construction. **(a-d)** Execution times until successful construction of matrices **Z**_*s*_, **Z**_*t*_, **W**_*y*_ and **W**_*r*_ (upper row), and number of neurons in the retrieved network (lower row) for the graphs analysed in Fig. 4. Median *±* [25th, 75th] percentiles are shown.

**Figure 6.**
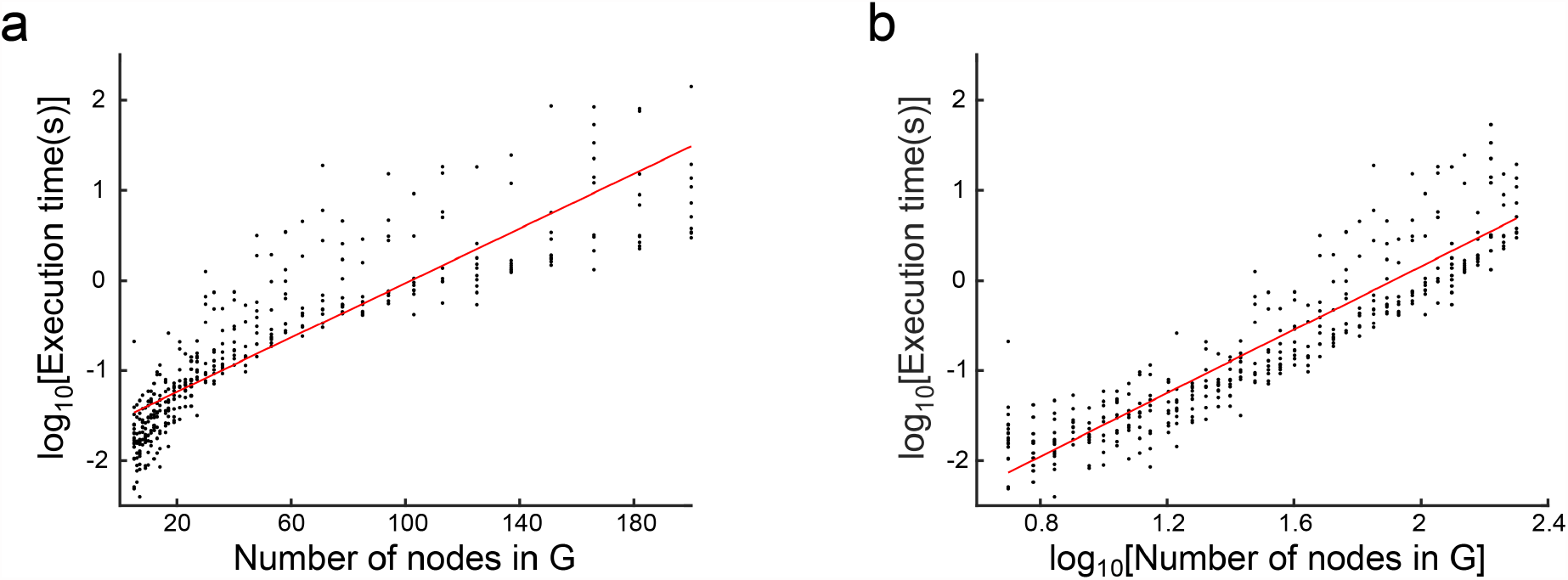
Polynomial execution time to attain delta consistency. **(a)** Execution time until consistency (*T*_*c*_) as a function of number of nodes (*N*_*v*_) in *G* (before expansion), is not well fitted by an exponential: log_10_*T*_*c*_(*N*_*v*_) = *a N*_*v*_ + log_10_*b, a* = 0.015, CI = (0.014, 0016), log_10_*b* = −1.54, CI = (−1.60, −1.48), d.f. = 398, R− squared = 0.77. **(b)** Execution time is well fitted by a power function: log_10_*T*_*c*_(*N*_*v*_) = *a* log_10_*T*_*c*_(*N*_*v*_) + log_10_*b, a* = 1.76, CI = (1.68, 1.84), log_10_*b* = −3.37, *CI* = (−3.49, −3.24), R squared = 0.82, d.f = 398. Execution times were computed for random graphs, with 3 stimuli and between 5 and 200 nodes, in logarithmic scale. A total of 10 graphs for each *N*_*v*_ value were constructed.

### Uncovering general dependencies between structure and function

We employed gFTP as a tool to study the multiple dependencies between neural dynamics and its underlying connectivity. To that end we employed a genetic algorithm (GA)[15] to optimize several measures computed on the transition graph and the synaptic weight matrix. We considered commonly employed graph theoretical measures, like the clustering coefficient of the transition graph, or the reciprocity of the recurrent synaptic weight matrix. Moreover, we computed the mean normalized mutual information between a target node and the stimulus that leads to it (abbreviated as “mutual information” (*I*)). We defined a population of transition graphs, and a mutation function that introduced variability by permuting two target nodes, or replacing one target node with another. We ran several experiments, which differed in which measure was maximized, although all measures were computed. Then, we computed the Pearson correlation coefficient (CC) between each pair of measures along the evolutionary process (e.g. Fig. 7a,b). In this way we expected to uncover how constraining (maximizing) one feature, either related to network dynamic or connectivity, impacts on the correlational structure of all other measures. On average, measures could be grouped together into two clusters, each defined by positive intra-cluster and negative inter-cluster CC (Fig. 7c): Cluster 1, composed of *N*_*v*_, *N*_*neu*_, *I*, and *Q* (modularity), and Cluster 2, composed of reciprocity (*r*), reciprocity of absolute synaptic weights (*r*_*abs*_), variability in average outer strength of the synaptic matrix (*σ*_*out*_), and the clustering coefficient of *G*_*cons*_ (*c*). The positive CC between *N*_*neu*_ and *N*_*v*_ in Cluster 1 is expected, since more neurons are required to generate enough population states. Also, the positive CC between *N*_*v*_ and *I* can be explained as an effect exerted by nodes added during graph expansion: new nodes only encode one stimulus when they are first added to the graph, thus contributing to a higher mutual information between node and stimulus. In Cluster 2, the positive CC between modularity and the clustering coefficient was also expected. Modularity was the only measure that showed a tendency for positive CC in most cases, even when the clustering coefficient showed negative CC. Reciprocity and absolute reciprocity were positively correlated with modularity and clustering coefficient, revealing an interesting relationship between a structural feature and a dynamical feature. Higher than chance reciprocity has been found experimentally[16], and it has been connected to optimal memory storage[17] and stimuli sequence encoding[13]. Results in Fig. 7 suggest that it may also subserve modular dynamic in general, which in turn can be related to specialization of function. The correlations analysed so far are clearly distinguished in the mean correlation matrix. A similar tendency can be recognized in the correlation matrix computed separately for each optimized measure, albeit with some departures from the average. To assess the immediate impact that an optimized measure has upon the others, we computed a correlation between two vectors of CCc: one vector collects the CCs between measure *i* and measure *j*, when measure *i* was optimized; the other vector collects the CCs between measure *i* and measure *j*, when measure *j* was optimized. These two vectors were positively correlated (Fig. 8a,b), meaning that there is a tendency to find the same dependences between measures regardless of which one was optimized. Even so, the unexplained variance was considerable. All in all, this suggests that the correlational structure between dynamical and structural measures has one component, which does not depend on the optimization process, and another component, which may depend on which measure is optimized, and even the entire optimization history.

**Figure 7.**
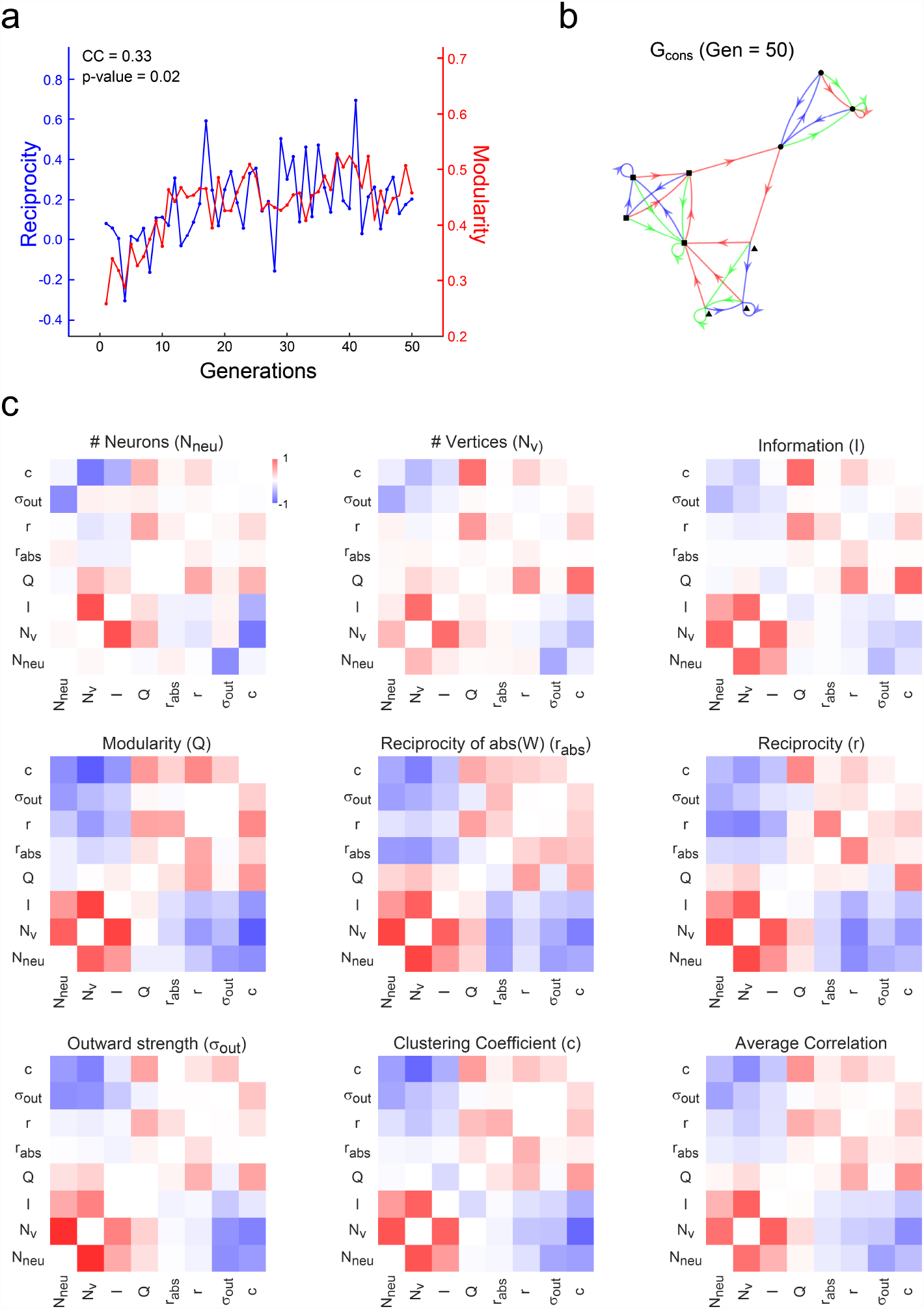
Correlational structure of dynamic and connectivity features in optimized networks. (a) An example of the evolutionary process. Reciprocity (*r*) and modularity (*Q*) are shown, computed for the elite individual in each generation of a 50 generations evolutionary process. Modularity was the fitness function in this case. Pearson correlation coefficient (CC) between both measures is shown, together with its p-value **(b)** Transition graph *G*_*cons*_ obtained with gFTP from the elite individual in the last generation of the evolutionary process shown in (a). Nodes plotted with different shapes (circle, square, triangle) indicating the three modules that maximized modularity. *Q* = 0.46, *r* = 0.19 for this graph and its associated network, respectively. **(c)** Correlation matrix between 8 measures that quantify distinct aspect of network dynamics and connectivity. Measures were always computed on the elite individual, obtaining one matrix for each repetition and for each fitness function. Each panel shows the correlation matrix, averaged across repetitions, for the fitness function indicated in the panel title. The matrix shown in the last panel is the average over all the other matrices. Color bar in first panel shows color scale for all panels (pure red for CC = 1, and pure blue for CC = -1).

**Figure 8.**
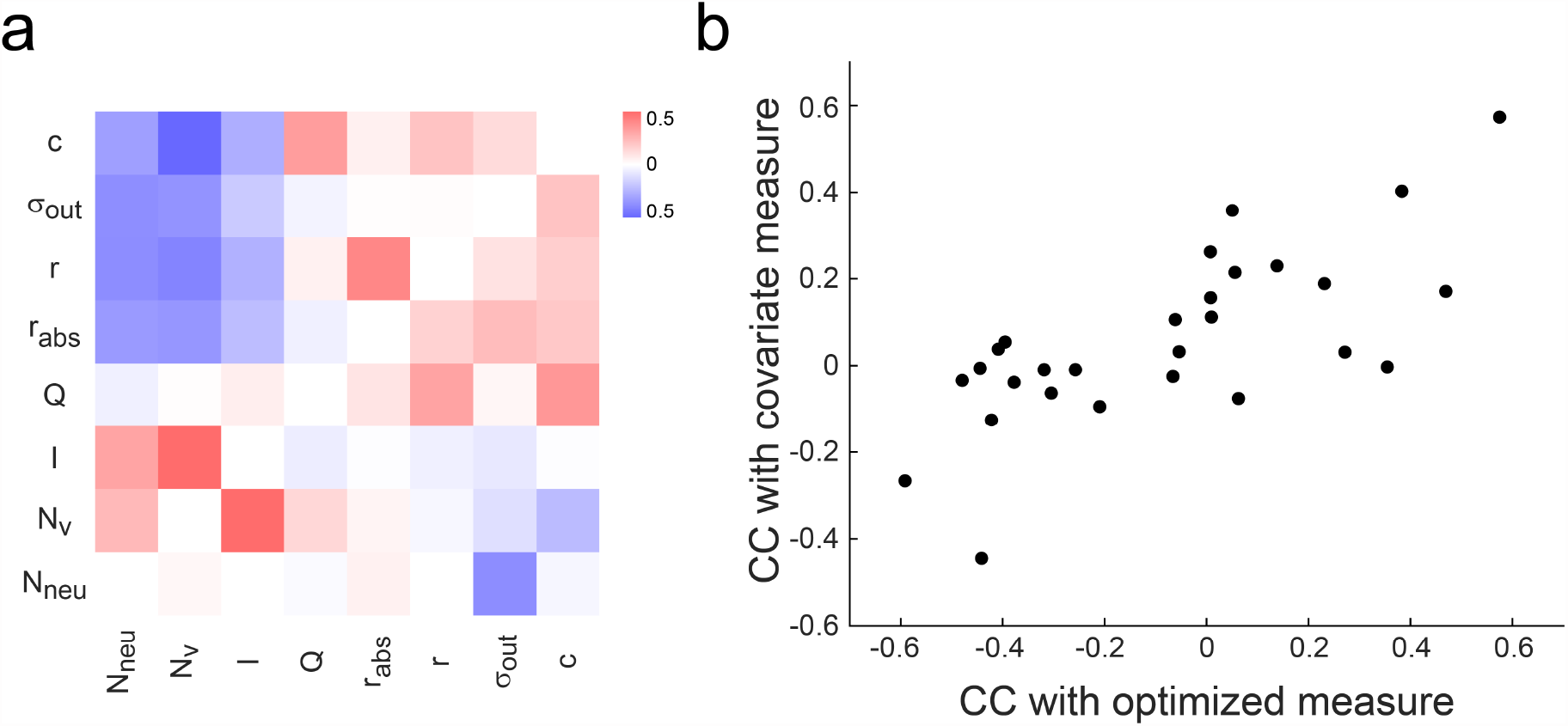
Dependence of correlation structure on the optimized measure. **(a)** Matrix showing correlations between measures along the evolutionary process. Row labels indicate the measure employed as fitness function, and collect correlation coefficients (CC) between the optimized measure and the other measures. A symmetric matrix would imply that correlations are completely independent of which measure of the correlated pair of measures was the optimized one. (b) Entry (*row, col*) in matrix shown in (a), plotted in the *x* axis, against entry (*col, row*) in the *y* axis. Spearman correlation *ρ* = 0.69, *p* = 8.10^−5^. A symmetric matrix in (a) would result in points lying in a line of slope = 1.

### Assessing dependencies between dynamic, connectivity, and the algorithm instantiated by the network

We sought to understand how two ways of solving the same task (two different algorithms) constrain the connectivity and dynamic of the network that solves it. To that end, we considered a stimulus discrimination task with two possible stimulus-response mapping, indicated by a context cue (Fig. 9a). The cue is presented first, followed by the stimulus to discriminate. Once the stimulus is presented, the network has all the information to execute the correct response, and therefore all information encoded in the neural population regarding context and stimulus can be discarded. We call this case Algorithm 1 (A1, Fig. 9b). Nevertheless, context and stimulus information could be retained, for purposes other than solving the task at hand, as has been observed experimentally[18]. This is Algorithm 2 (A2, Fig. 9c). We constructed networks for each algorithm, and measured the number of nodes in the consistent graph, mutual information between stimulus and node, the number of neurons in the constructed network, and its reciprocity. We also assessed the impact of redundancy in network states. Redundancy was accomplished by expanding the initial transition graph with new nodes that encoded no more information than the original nodes (similar to expansion when constructing a consistent graph, see *Methods*). Algorithms 1 and 2 could be sorted out based on these measures, despite the small differences between their transition graphs (Fig. 9d-g). Higher redundancy led to higher *N*_*v*_ and *I* in the consistent graphs, and higher number of neurons and lower reciprocity in the networks, for both algorithms. This is consistent with results in Fig 7. However, with matching redundancy levels, Algorithm 1 showed lower *N*_*v*_ than Algorithm 2, but higher *I*, contrary to the positive correlation found in Fig. 7. Networks executing Algorithm 1 had fewer neurons than networks executing Algorithm 2, and less reciprocity, which is also in opposition to the negative correlation these two measures exhibited in Fig. 7. If these results generalize, they would suggest that mixed selectivity to stimuli (lower *I*) and higher reciprocity are among the distinctive traits of networks that instantiate more complex algorithms. They would also imply that the dynamic and connectivity measures here considered depart from the main dependencies observed in Fig 7 when specific transition graphs are analysed.

**Figure 9.**
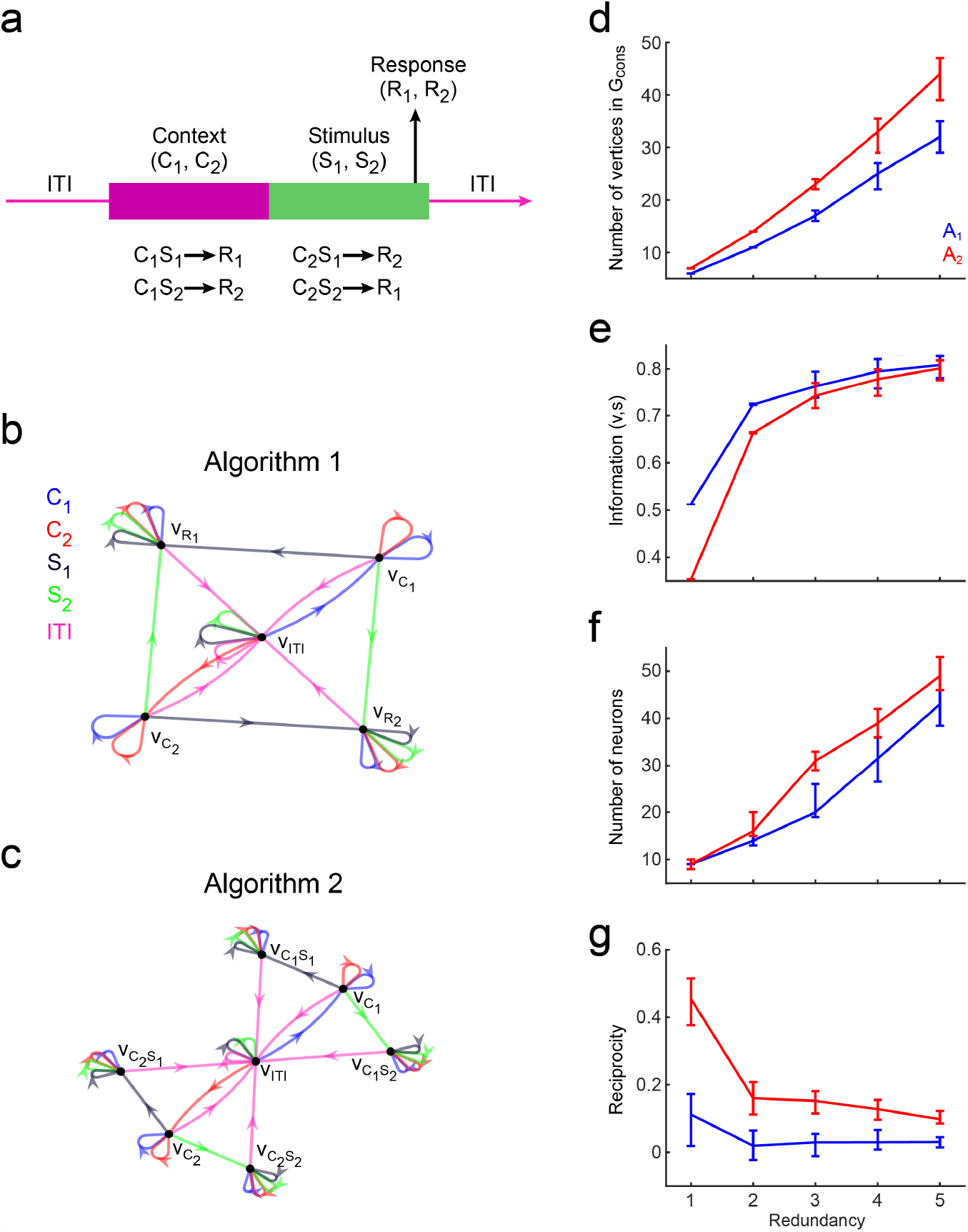
Effect exerted by the instantiated algorithm on the network dynamic and connectivity. **(a)** Context-dependent discrimination task. A trial starts with the presentation of a context cue, indicating the correct stimulus-response mapping for the trial, followed by the stimulus to discriminate, time at which the agent is expected to execute the correct response. **(b)** Transition graph for Algorithm 1 (A1). nodes 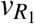 and 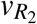 encode the correct response, which deppends on context and stimulus. Once the system is in one if these states, it is impossible to recover the stimulus and context that led to the decision. **(c)** Transition graph for Algorithm 2 (A2). Similar to A1, but it has nodes 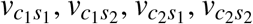, wich simultaneously encode stimulus and context, and therefore convey the necessary information to trigger the corresct response (*R*_1_ with 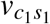 and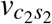, and *R*_2_ with 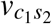 and 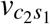). Self loops render states refractary to stimuli that are out of place, like a context cue during stimulus presentation, or a second presentation of a stimulus when the stimulus was already presented and encoded. **(d-g)** Quantification of differences in dynamic and connectivity features between networks instantiating A1 or A2, with increasing levels of redundancy. Redundancy shown in the x axis indicates how many times the number of nodes in graphs shown in (c-d) was multiplied by adding redundant nodes. Measures were computed on the *G*_*cons*_ and synaptic weight matrices obtained through gFTP. Median *±* [25th, 75th] percentiles are shown, computed on 100 graphs and their respective weight matrices, per redundancy level.

## 4 Discussion

We have introduced an algorithm that takes a target dynamic as input and returns the connectivity matrix of a neural network that follows the target dynamic. This approach departs from other model building approaches in several ways. In bottom up approaches, a model is a specific formalization of a series of proposed mechanisms. Its dynamic is a complex emergent of these mechanisms, and in most cases it cannot be specified in advance. Although it is possible, for some models, to prove analytically some of their properties (like in [19, 20]), this is mostly done at the cost of many simplifications. Complex dynamics that can instantiate specific algorithms are difficult for analytic treatment. On the other hand, normative approaches constrain dynamics to the subsets of dynamics that minimize the loss function, but neural network models have enough degrees of freedom to find very disparate solutions. While a multiplicity of solutions is appealing in itself, as a way of exploring novel dynamics, it also makes it problematic to go from specific solutions to general conclusions regarding mechanisms. In this sense, our algorithm brings a new angle to the way we build neural networks as models of brain function. The gFTP algorithm starts by defining the dynamic that will be unfolded by the network, specified as a transition graph in which nodes are network activation states, and arcs are transitions between states, triggered by stimuli. Then, the algorithm constructs an auxiliary graph (the *D* graph). If this graph has a cycle, then the transition graph cannot be realizable by a neural network. The algorithm proceeds by expanding the original transition graph, adding nodes that break cycles in graph *D*, without loosing any of the information encoded in each network state. Eventually, all cycles are removed, and the final transition graph is realizable. Thus, gFTP provides a high level of control over the network dynamics. In turn, this higher control can be exploited in several ways:

1. We can, for instance, construct networks which exhibit dynamics that have been proposed theoretically, or found experimentally, like the discrete attractor dynamic proposed to subserve working memory, or the toroid dynamic proposed to subserve spatial coding. Also, we can define dynamics by means of the algorithms that they instantiates, either at the level of cognition or behaviour. This aspect is particularly specific to our approach, since normative methods are not well suited for specifying exactly how the network solves a task. This makes gFTP particularly useful for studying the interdependencies between dynamic and the algorithms it implements, the information encoded in the activation states, and network connectivity. We may define different algorithms that solve the same task and then compare the expanded graphs, or the networks obtained, and draw conclusion about how the differences in the algorithms translate into differences in dynamics or network connectivity. If it happens that gFTP finds a graph to be inconsistent, then it means that the instantiated dynamic cannot, as such, be realizable in a neural network. This fact is in itself useful, because we can study which features of the dynamic are essential to a given algorithm, and what features are there because of the constrains imposed by the fact that dynamic has to be carried out by the neuronal machinery.
2. For each transition graph that is inconsistent there are many different ways to expand it into a consistent one. We can then run gFTP multiple times, sampling from the distribution of expanded graphs that come from the same initial graph, and study how different the expanded graphs can be.
3. Once we have a consistent graph, we can generate matrices **Z** in a way that they all share some target activation statistics (e.g. same mean population firing rate and correlation). Then, we can assess how the dynamics/algorithms constrain the variability of those statistics.
4. For each dynamic, and matrix **Z**, we can generate a sample of synaptic matrices that instantiate the same dynamic with the same activation states, and assess how dynamics and firing statistics impact on network connectivity.

In sum, the gFTP algorithm provides a high level of control over key aspects of analysis, like network function, dynamics and firing statistics, allowing to disentangle their interactions by carrying out controlled numerical experiments.

Our results with random graphs showed a time complexity well fitted by a power function of the number of nodes, with exponent ∼1.76, for the operations of consistency assessment and expansion. This probably stems from the fact that the main subroutine for these operations is Depth First Search, which has linear time complexity on the number of nodes and arcs (which becomes quadratic at most, in the number of nodes, in the case of a fully connected graph). On the other hand, to construct **Z**_*t*_ our current implementation explores the tree of possible assignments for each neuron in each network state, accelerating the process by computing deltas and filling assignments that are completely specified by the deltas and *z* values already defined. We also used the heuristic of assigning *z* value first to nodes with higher degree in graph *G*, for these are expected to be part of cycles with higher probability. This heuristic helped in reducing execution time during construction of matrices **Z**_*t*_. Yet, the algorithm would benefit from other ways of assigning *z* values. Studying the structure of graph *D* and cycle formation will certainly help. Also, optimization methods like genetic algorithms could be good options to evaluate.

Once matrices **Z**_*t*_ were constructed, we found matrices **W** by employing the accelerated perceptron algorithm. We decided to add neurons to the network in multiples of the number of nodes in the transition graph. The reason for this criterion is that, for matrix **W** to exist, matrix **U** must have the same linear combinations that matrix **C**, and H(**U**) = **Z**_*t*_. Not all matrices **Z**_*s*_ and **Z**_*t*_ allow the existence of a matrix **U** that fulfills this two conditions. Yet, the probability of finding one viable matrix **U** is expected to be higher if the number of linear combinations in **C** is low. This can be achieved by adding neurons to the network, with random values of *z*, since the probability of a linear combination between rows of **Z**_*s*_ is expected to drop with the number of its columns. Adding neurons in multiples is also convenient in that the number of calls to the accelerated perceptron is reduced: too few neurons would certainly not lead to a solvable system, and the accelerated perceptron will have to reach the maximum number of allowed iterations in order to stop and try with a higher number of neurons, thus increasing execution time. In our experiments, a number of neurons approximately matching the number of nodes were enough to define a solvable system of linear separation problems.

One limitation of our method is that it is restricted to a simple neuron model. The binary neuron is not a proper model of membrane potential, because preactivations only depend on the neuron inputs, and not on preactivations in the previous time step. Therefore, the model cannot give spike trains with realistic time intervals. A network of binary neurons is suitable, nonetheless, to explain dependences between dynamic, firing statistics, and connectivity which do not heavily depend on a detailed model of the membrane potential dynamic[17, 21]. Another limitation is that, for constructing matrix **Z**_*t*_, we rely on a backtracking approach, which can be inefficient. Better methods could be developed in the future for this task, as we have discussed above. Also, matrix **Z**_*t*_ could be generated only to be close to consistent, and **W** only leading to an approximation of **Z**_*t*_. In this case, the obtained network will not follow the target dynamic exactly. However, it is expected that, with enough neurons and redundancy in **Z**_*t*_, the approximate network will follow a dynamic that is close enough to the target. It could also be possible to use the network found with gFTP as a good starting point for further optimization with another algorithm.

As we have already stated, the gFTP algorithm is suitable for studying the interdependence between network parameters and its function, by generating a sample of networks that share a given target property. Recent works have address similar goals. In Brennan *et al*.[22] a method termed LOOPER was introduced, in which a Markov process is constructed from recorded neural activity, with transition probabilities modelled as an asymmetric diffusion process, such that the global dynamic respects convergences and divergences observed in the data. The method relates to ours in that it allows to construct a model of an arbitrary dynamical system. This is accomplished in our algorithm in the transition graph, in which convergences and divergences are what differentiate one graph from another. LOOPER has the advantage of finding a model directly from data, while the use of a binary neuron model in gFTP makes it more difficult to translate recorded firing activity into a transition graph. However, and in contrast to our method, LOOPER never constructs a neural network model, so there is no explicit way of linking the low dimensional dynamic recovered in the model with any underlying network connectivity. On the other hand, in the work of Brittner *et al*.[23] and Gonçalvez *et al*.[24] a deep neural network was used to approximate a probability density function over parameters of a target neural network, such that a target mean over an emergent property is satisfied. The method proposed by Brittner *et al*., called Emergent Property Inference (EPI), is very versatile, in that the target model only has to be differentiable, but it does not give control over specific dynamics, because different dynamics can show the same emergent property. In Gonçalvez *et al*. the method called Sequential Neural Posterior Estimation (SNPE) is introduced, in which a distribution over parameters is fitted to the observed data, or data summary features computed from the data. In this case the distribution is fitted through data collected by simulation, which gives the possibility of assessing non differentiable models. The computational cost of both methods is an issue: when tested to find distributions of low rank recurrent networks with up to 1000 parameters, constrained to have synaptic weight matrices of given eigenvalue, SNPE failed to converge with the biggest networks, while for EPI it took up to 5 hours of training. While impressive, this is a fitting task with constrains on the synaptic weight matrix, which is expected to be easier than with constrains on network activity. Although gFTP generates one network at a time, each generation is fast, hence a big population of networks can be gathered. Also, it is guaranteed that all networks will have the desired dynamic. Beside, gFTP can always be adapted to generate networks that only approximate the target dynamic, as discussed above. In conclusion, we have introduced a systematic way of constructing transition graphs that are realizable by a neural network of binary neurons, allowing to test hypothesis regarding function, dynamic and connectivity in ways not matched by other methods. We believe the gFTP algorithm is going to be a valuable addition to the toolbox of the theorist.

## 5 Methods

### Constructing matrices Z*s* and Z*t* with backtracking

To ensure that all nodes are differentiated, we define neurons so that their output is different for at least a pair of nodes. Thus, we choose two nodes (*v*_1_ and *v*_2_) that are going to be differentiated. First, the algorithm assigns value 1 to node *v*_1_ and 0 to *v*_2_. Next, values 1 or 0 are assigned to each one of the remaining nodes, one at a time, following a list *L*_*u*_ of undefined nodes sorted in decreasing in-degree order. The first element in this list is set as the active element, the one that is going to be defined. Any node that is set as active is registered in a list of active elements *L*_*a*_. Each new assignment may define a new delta, so delta consistency must by checked. If consistency is sustained, new node assignments are made as necessary, according to the *z* and delta values defined so far (propagation of values according to delta consistency). Then, consistency is checked again. If sustained, the first non defined node in *L*_*u*_ is set as the active element. The process continues with the new active element until all nodes have *z* value. If delta consistency is violated, another *z* value is tried for the active element (1 if 0 was already tried, 0 if 1 was already tried). If neither of the assignments was successful, the algorithm makes a step back by searching for the last successfully assigned active element in *L*_*a*_, making it active, and erasing from the list all elements after it. All nodes that were defined after the active element are set as undefined. Assigned values are removed from the list of viable values for that node, but all values are restored after a step back, for all the following nodes in *L*_*u*_, after the active one. If all *z* values were tried for the first node in *L*_*u*_, without success, then the algorithm stops with no **z** as output (there is no node left to step back), and hence the transition graph is delta inconsistent. A graph that was previously expanded to obtain a consistent graph *G*_*cons*_ always leads to a fully assigned **z**_*neu*_ for any pair of differentiated nodes.

Once we get vector **z**_*neu*_, we identify all nodes that were differentiated (which can be more than just *v*_1_ and *v*_2_), and remove them from the list of pairs of nodes to differentiate. If all pairs of nodes were already differentiated, then each node has one vector **z**_*v*_ that encodes that node. The set of all **z**_*v*_ are arranged to construct matrices **Z**_*s*_ and **Z**_*t*_. These two matrices define *N*_*neu*_ linear separation problems (one for each neuron), where the *N*_*tran*_ column vectors in **Z**_*s*_ are the points to separate, and the *N*_*tran*_ ones and zeros of each row in **Z**_*t*_ are the classes. For each neuron we find its input weights by fitting a perceptron through the accelerated perceptron learning algorithm (algorithm No. 2 in [14]). If at least one neuron is encountered for which the separation problem cannot be perfectly solved, we assume that at least one linear combination in **Z**_*s*_ cannot be reproduced in **U**. In this case, new matrices **Z**_*s*_ and **Z**_*t*_ are generated and concatenated to the previous ones, to increase the number of neurons and reduce the number of linear combinations in **Z**_*s*_. This process is repeated until each neuron can be perfectly solved.

In practice, we frequently observed cases where all pairs of nodes were differentiated but the accelerated perceptron failed to find a solution, meaning that more neurons were required. Therefore, each round of **Z**_*s*_ and **Z**_*t*_ matrices were constructed by adding neurons until all pairs of nodes were differentiated *and* the total number of neurons surpassed the number of nodes in *G*_*cons*_.

### Transition graphs analysed

#### Random transition graphs

We set a number of nodes and stimuli. For each node, we defined its target nodes reached through each stimulus. Nodes were numbered, and target nodes were taken at random, with replacement, from the list of nodes that went from 2 nodes before the source node, to 2 nodes after the source node, avoiding self-targeting. Therefore, transitions were at random, but constrained to a vicinity around the source.

#### 2-dimensional continuous-like attractor

We defined a discrete 2-dimensional space, square in shape, of a certain integer side length. There was one node for each discrete position in this space (i.e. 9 positions and nodes for a 3×3 square arena). Parallel sides were glued together, forming a torus. We defined 5 stimuli, 4 for vertical or horizontal displacements, and a 5th for stillness. Therefore, the transition graph instantiated a discretized version of the dynamic of a continuous attractor model, thoroughly studied in the context of spatial coding [11].

#### Discrete attractors

we defined a number of nodes, and a number of stimuli, which was the same than the number of discrete attractors (each stimulus had an associated attractor node, and that stimulus led to its associated attractor). Then, we assigned to each node a random position in a 2-dimensional space, and defined a graph in which each node was connected to its closest 6 nodes, measured in Euclidean distance between positions. Finally, we found the shortest path from each node to each of the attractor nodes. We constructed graph *G* by making each stimulus trigger a transition to the first node in the shortest path pointing to the attractor associated with that stimulus. Since some nodes could end up being source without being target, for each one of these sourceless nodes we introduced one extra node, that went to the sourceless node through one stimulus, and to itself through the remaining stimuli.

### Context dependent discrimination

We considered a task in which the agent must choose one out of two possible responses when presented with one out of two possible stimuli. There were two rules (or mappings) by which stimulus and response might be matched: (*S*_1_, *R*_1_), (*S*_2_. *R*_2_), or (*S*_1_, *R*_2_) (*S*_2_, *R*_1_). In each trial the rule is chosen at random, and indicated through a context cue (*C*_1_ or *C*_2_), which is presented immediately before the stimulus to discriminate. The response is observed during stimulus presentation. Trials are separated by a inter-trial interval (ITI). We considered a recurrent neural network that receives the context cue and stimulus as input. It also receives a specific input that represents the ITI, and puts the network into a basal state. The context cue pushes the network to a context coding state (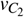 and 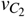). From this state, the network is sensitive to the stimulus, which triggers a transition to the correct response coding state. We did not model a dedicated network to execute the response, and instead we only cared that the correct response was decodable from the population state. We considered two possible algorithms for this task. In Algorithm 1 stimuli triggers transitions from *v*_*C*_ to the population state that encodes the correct response. This state can be reached through stimulus *S*_1_ or *S*_2_, depending on context, implying that states 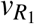 and 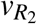 do not encode context nor stimulus *per se*. Therefore, the information regarding context and stimulus is lost when the response is executed. On the contrary, in Algorithm 2, the population state that encodes the correct response is different depending on the context state and the stimulus presented. This causes the states that encode the correct response to also encode the context and the stimulus presented. All the information is preserved.

### Optimization with a genetic algorithm

We constructed a population of 30 individual. Each individual is a transition graph *G*, possibly not realizable, with *N*_*v*_ = 5 and number of stimuli *N*_*s*_ = 3. For each *G* a graph *G*_*cons*_ is constructed with gFTP, together with the synaptic weights of the network that instantiates *G*_*cons*_. Next, the fitness of each individual is computed, by applying the fitness function to its *G*_*cons*_ or synaptic weight matrix of that individual. The fitness function is chosen from a set of measures, listed below. Then, individuals are selected with replacement with probability proportional to their fitness. A mutation operator introduces variability to each selected individual by applying one of two equally likely modifications to *G*: permutation of the target node of two different transitions, or replacement of the target node of a transition by another node. In the last case, only nodes that appear more than once as targets are considered to be replaced, to avoid the desappearance of any node. Therefore, all graphs will remain having *N*_*v*_ nodes throughout the evolutionary process. The new population is composed of the mutated individuals, plus an unaltered copy of the best individual in the last generation (called the elite individual). We were interested in finding how maximising a given feature of the transition graph or the synaptic weight matrix affects other dynamical or connectivity features. To do this, we computed a series of measures on the elite, and kept track of them along the evolutionary process. To obtain the data summarised in Fig. 7, we conducted the evolutionary process 20 times, during 50 generations each, computed correlations between the tracked measures, and average them over the 20 repetitions.

### Measures computed over transition graphs and synaptic weight matrices

The measures analysed and selected as fitness function were:

- *N*_*v*_: the number of nodes in the consistent graph obtained through gFTP applied to a given individual in the population.
- Clustering coefficient (*c*): Defined as the average computed over all per-node clustering coefficients. The clustering coefficient of a node is the number of triangles the node is part of, divided by the number of all possible triangles there could be for that node[25].
- Modularity (*Q*): the number of arcs within modules minus the expected number of arcs in a random graph with matching degree distribution. We employed the Leicht and Newman algorithm[26] to partition nodes into modules that maximise *Q*.
- *I*: the average information encoded in a network state about the stimulus that leads to it. Formally, the average normalized mutual information between a node taken as target and the stimuli that leads to it:

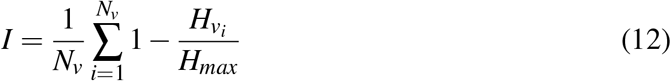

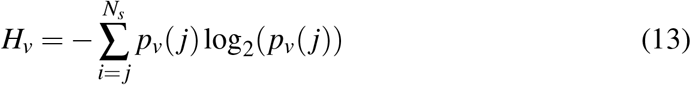

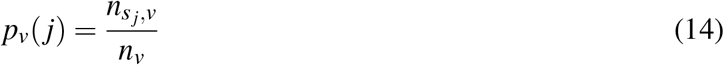

with *N*_*s*_ the number of stimuli, 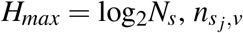 the number of arcs with label *s* _*j*_ that target *v*, and *n*_*v*_ the total number of arcs that target *v*.
- *N*_*neu*_: the number of neurons in the recurrent neural network.
- Reciprocity (*r*): The Spearman correlation coefficient computed between a vector collecting the weights from neuron *i* to neuron *j*, and a vector collecting the matching weights from neuron *j* to neuron *i*.
- Absolute reciprocity (*r*_*abs*_): Reciprocity computed over the unsigned recurrent synaptic weights.
- Standard deviation of out-strength (*σ*_*out*_): measures the variability on the average outward synaptic weights. We compute, for each node, the outward strength, defined as the mean over outward weights. Then, we computed the standard deviation over the outward strengths of all nodes.

The clustering coefficient and modularity were computed with the Brain Connectivity Toolbox[27]. Graph measures included all arcs in the graph, regardless of their labels.

Before computing any synaptic weight measure, we first divided the inward weight vector of each neuron by its Euclidean norm. This transformation leaves the neuron input-output mapping intact, and reduces weight variability accumulated during the accelerated perceptron learning process.

The minimum value for reciprocity and absolute reciprocity is -1, hence we added +1 to these measures when used as fitness function.

### Adding redundancy to *G*

Redundant codes are a hallmark property of robust systems, neural systems included[28, 29]. We introduced redundancy to a transition graph *G* by adding nodes that encode the same information than another preexisting node. Two nodes encode the same information if they are reached by the same sequence of stimuli and nodes. To accomplish this, we added new nodes, each one equivalent to a randomly chosen preexisting node, i.e. the new node leads to the same target node, and through the same stimuli, than its equivalent node. We say that equivalent nodes are members of the same equivalence set. Each node belongs to one equivalence set. There is one equivalence set for each node in the original graph, and they are mutually disjoint. Each time a new node is added, its target nodes are taken at random from a randomly chosen equivalence set. For results in Fig. 9 we added one new node for each equivalence set, and repeated this process *N*_*red*_ times, with *N*_*red*_ being the redundancy level.

## Acknowledgements

This research was supported by the Consejo Nacional de Investigaciones Científicas y Técnicas, Grant No. 11220200102316CO, Agencia Nacional de Promoción de la Investigación, el Desarrollo Tecnológico y la Innovación, Grant No. PICT-2021-I-A-00957, and Universidad de Buenos Aires.

## Appendix

Below we provide pseudocode for the gFTP algorithm and its key subfunctions, together with the following notes:

- The expression *x*(1) represents the first element of a vector or list, while *x*(*end*) is the last element.
- The expression [*x*_1_… *x*] stands for concatenation of *x*_1_… *x*_*n*_.
- The function DFS performs Depth First Search. When a cycle is encountered, the program stops and returns the arcs (*v*_*s*_, *v*_*t*_) that compose the cycle, and also a function *delta* that maps each arc with the *δ*_*i, j*_ that labels the arc.
- The graph *D*^*′*^ is like graph *D* but with its arcs flipped when there were antiparallel superimposed arcs. It is equal to *D* if no antiparallel superpositions were found.
- The set *V*_*di f*_ is the set of all pairs of nodes that are differentiated by the neuron (their assigned *z* values are different).
- The difference between two sets *A* and *B* is written *A*−*B*.
- In Algorithm 1: gFTP, *max_iter* is the maximum number of iterations executed by the accelerated perceptron algorithm while searching for a solution. It was set to 1000*N*_*tran*_.
- In Algorithm 3: Find_Superposition, when two sets in *P* are merged, the sets are removed from *P*, and a new set is added, that is the union of the two merged sets.
- In Algorithm 4: Make_Z_Target, the function *z*_*valid*_ takes a node as argument and returns a vector, specific for that node, which can be (1,0) or (0,1), chosen at random from a Bernoulli distribution. These vectors are defined once at the beginning of the algorithm, and are kept constant thereafter. The variable *firing_rate* is the parameter of the Bernoulli distribution. It was set to 0.5 in all experiments.

Algorithms were implemented in MATLAB R2021a (The MathWorks, Inc., Natick, MA, USA), running on a notebook computer with Intel™ i7-8550U CPU and 32 GB RAM.

### Algorithm 1 gFTP

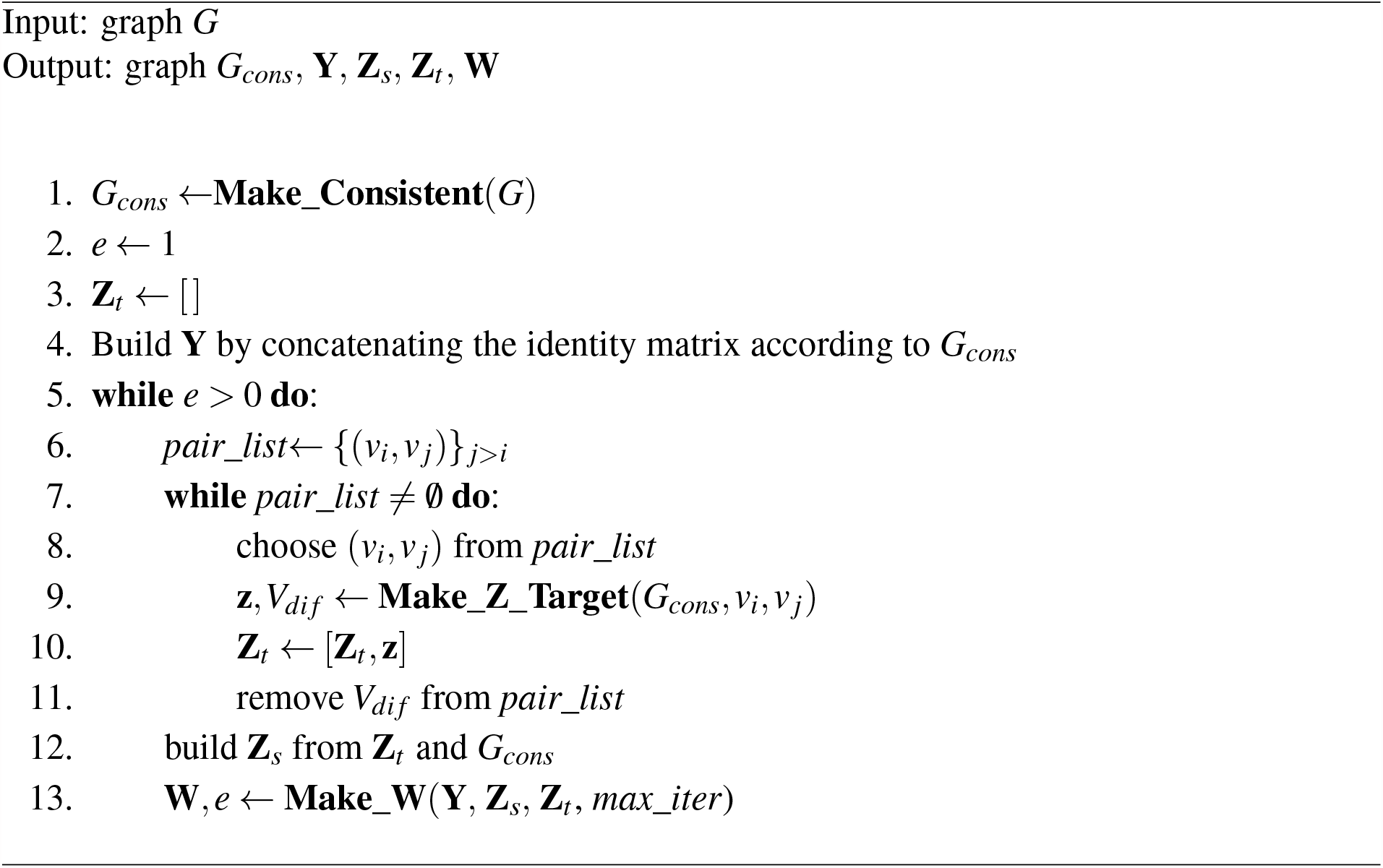

### Algorithm 2 Make_Consistent

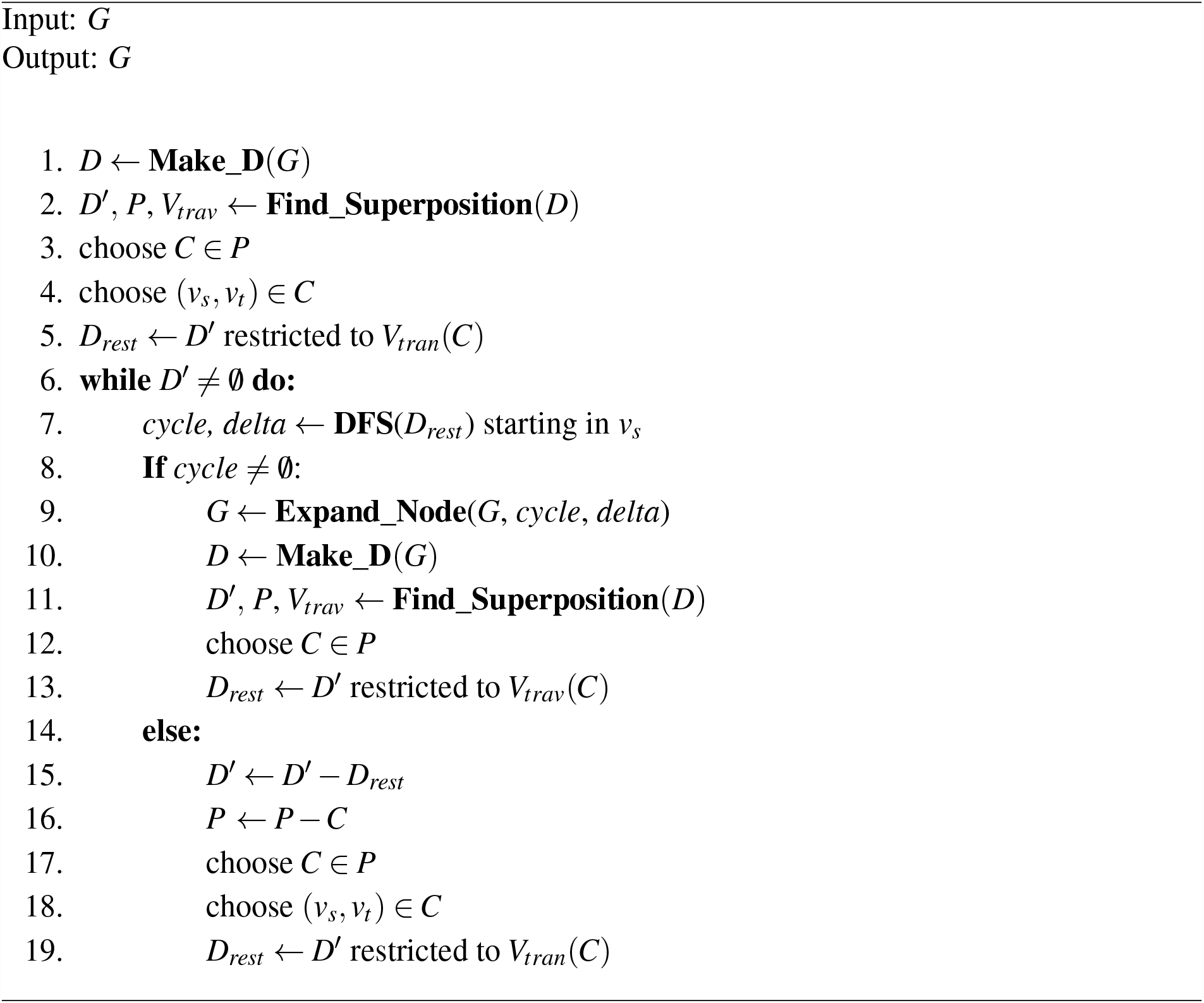

### Algorithm 3 Find_Superposition

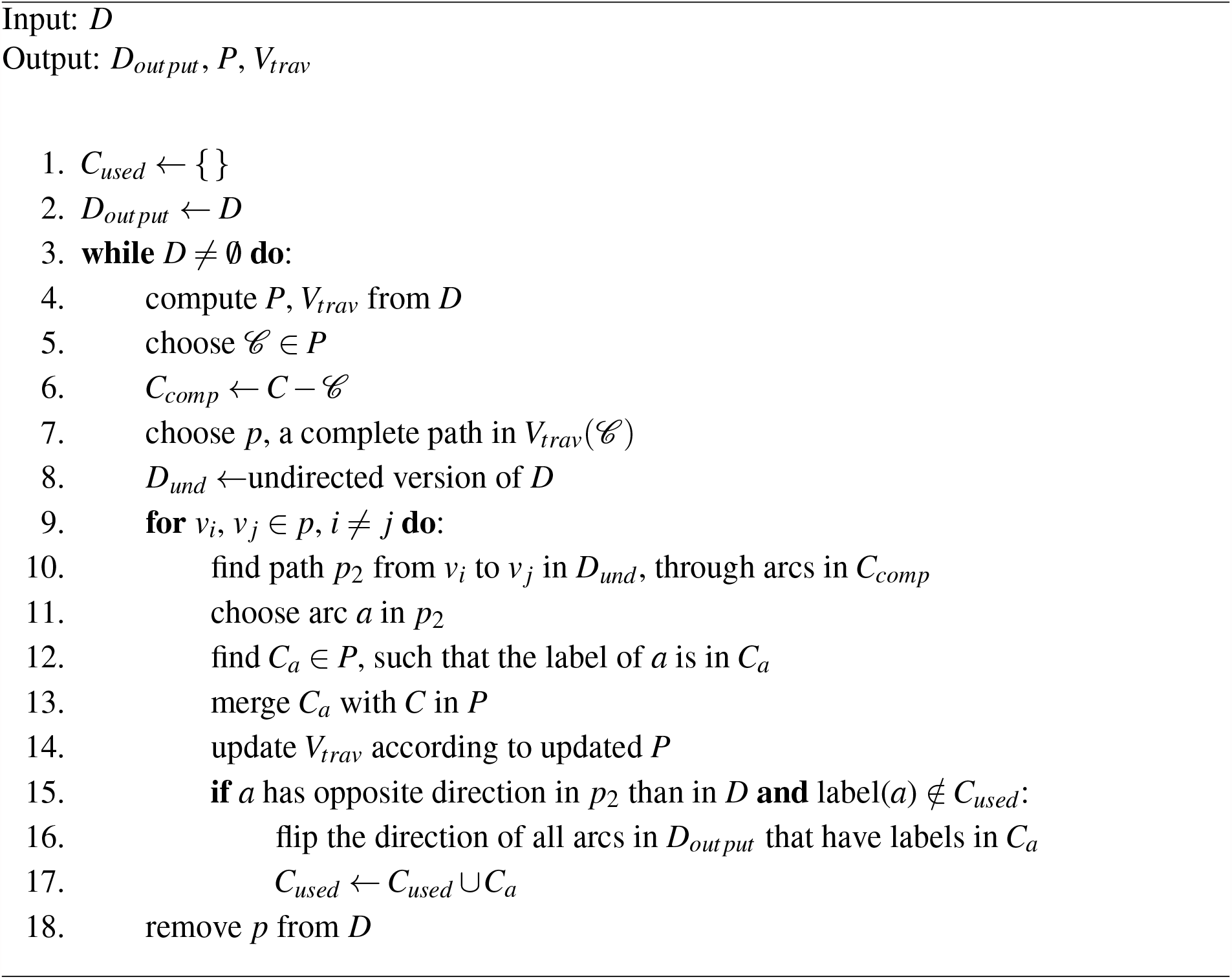

### Algorithm 4 Make_Z_Target

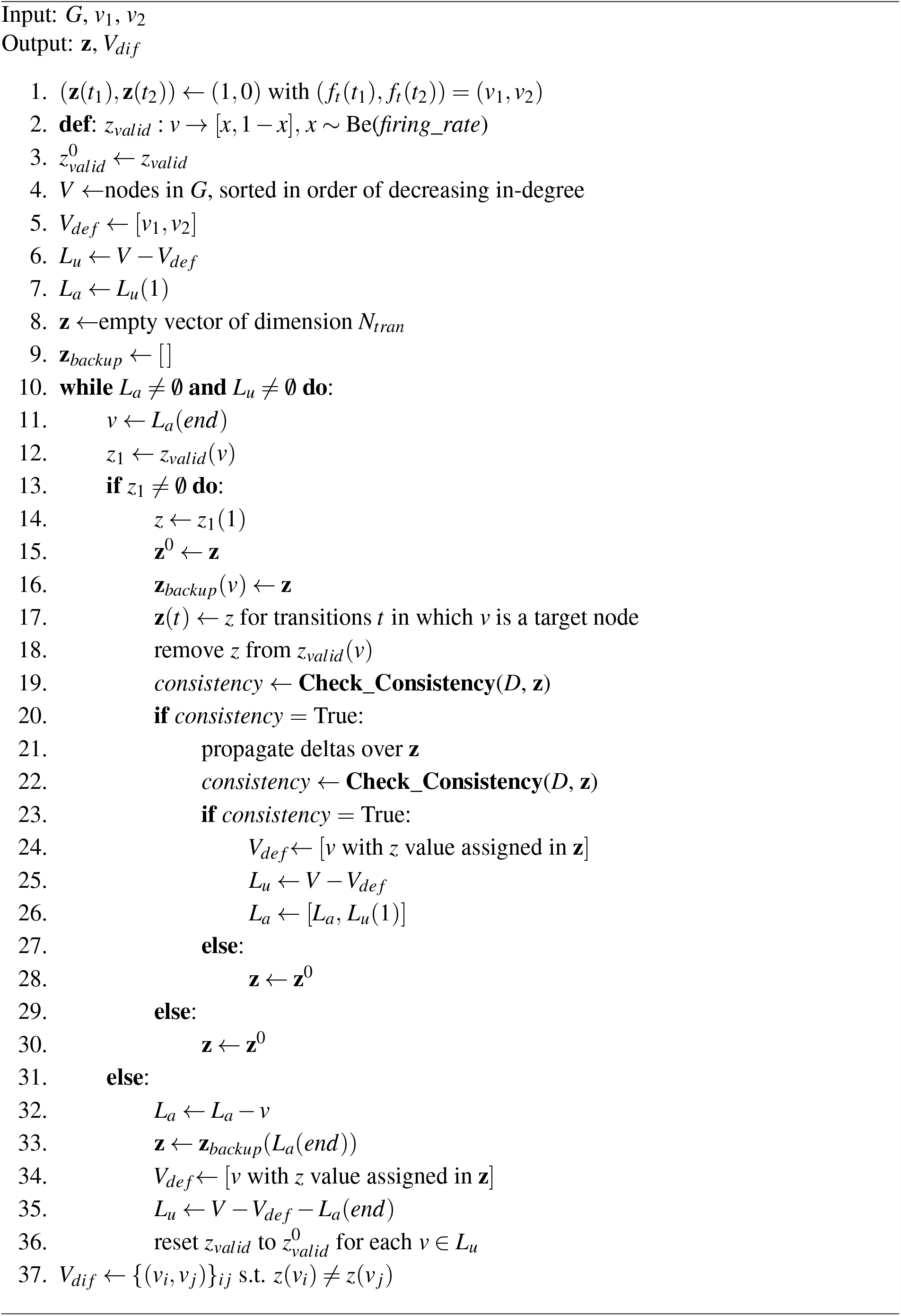

### Algorithm 5 Expand_Node

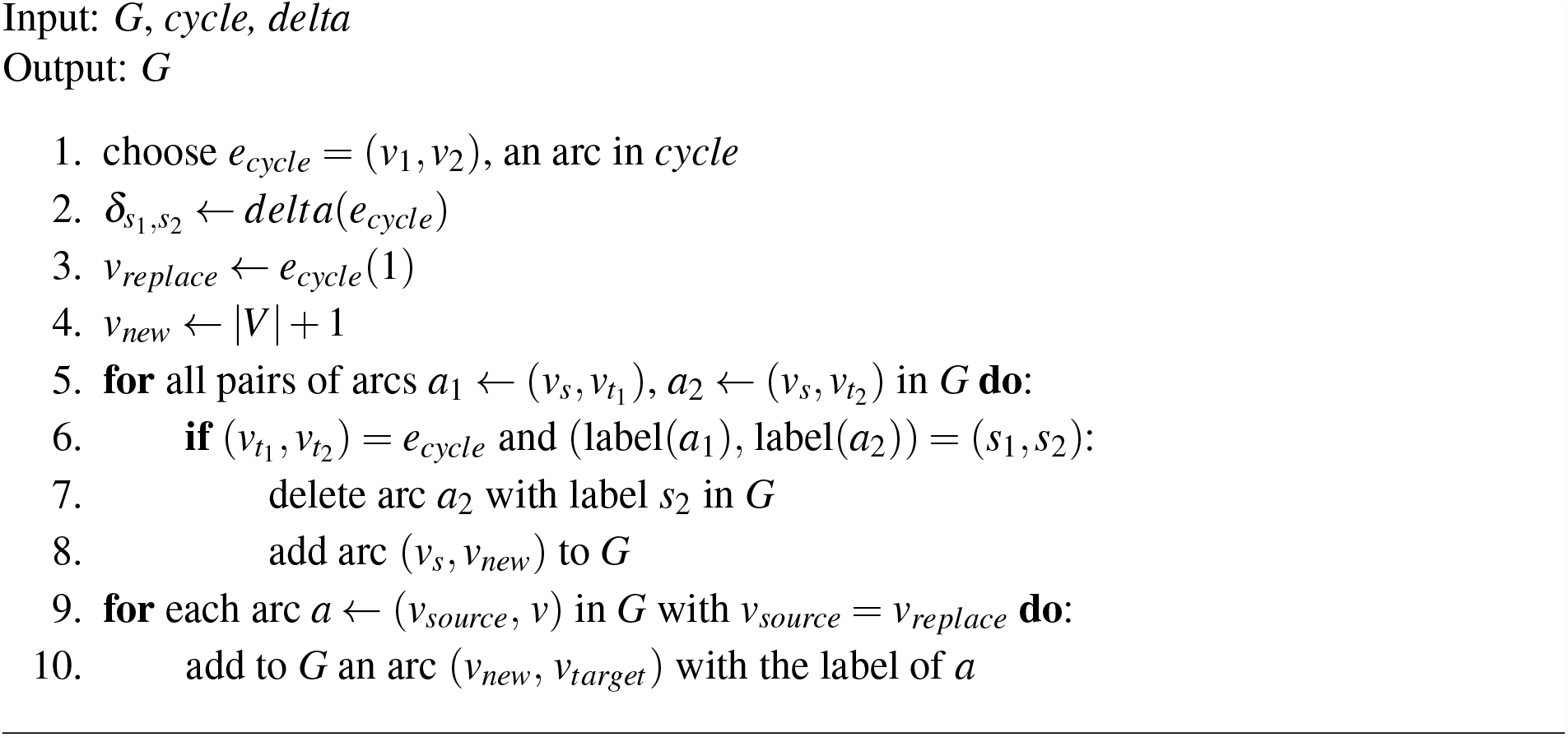

